# Discovery of novel quinoline papain-like protease inhibitors for COVID-19 through topology constrained molecular generative model

**DOI:** 10.1101/2024.09.07.611841

**Authors:** Yongzhi Lu, Ting Ran, Qi Yang, Guihua Zhang, Jiayi Chen, Peiqi Zhou, Wenqi Li, Minyuan Xu, Jielin Tang, Minxian Dai, Jinpeng Zhong, Hua Chen, Pan He, Anqi Zhou, Bao Xue, Jiyun Zhang, Kunzhong Wu, Xinyu Wu, Miru Tang, Xinwen Chen, Hongming Chen, Jinsai Shang

## Abstract

The rapid emergence of drug-resistant SARS-CoV-2 variants poses a persistent challenge to current antiviral strategies. Mutations in key viral targets, including RNA-dependent RNA polymerase (RdRp), main protease (3CL^pro^), and papain-like protease (PL^pro^), have been shown to markedly reduce the efficacy of approved therapeutics, highlighting the urgent need for next-generation antivirals capable of overcoming resistance. Here, we report the discovery of a novel class of PL^pro^ inhibitors through an AIDD strategy based on a topology-constrained molecular generative model (Tree-Invent) integrated with structure-guided optimization. Scaffold hopping from a previously reported lead enabled the identification of a quinoline-based chemical series with substantially improved metabolic stability and antiviral potency. Structure-guided optimization yielded compound GZNL-2016, which exhibited potent enzymatic inhibition of PL^pro^ (IC_50_ = 10.5 nM), robust antiviral activity against multiple SARS-CoV-2 variants, including Omicron BA.5 and XBB.1, and favorable pharmacokinetic properties following oral administration. Notably, GZNL-2016 retained substantial inhibitory activity against the clinically relevant drug-resistant mutant PL^pro^ E167K (IC_50_ = 480.2 nM; K_i_ = 439.3 nM), in contrast to previously reported inhibitors that exhibit markedly reduced potency. In a SARS-CoV-2 infection mouse model, oral administration of GZNL-2016 significantly reduced pulmonary viral titers, demonstrating in vivo antiviral efficacy. Collectively, this study establishes an AI-enabled strategy for rapid antiviral discovery and identifies GZNL-2016 as a promising lead compound to address the threat of coronavirus infections caused by drug-resistant mutant SARS-CoV-2 variants.

**Graphical abstract:** 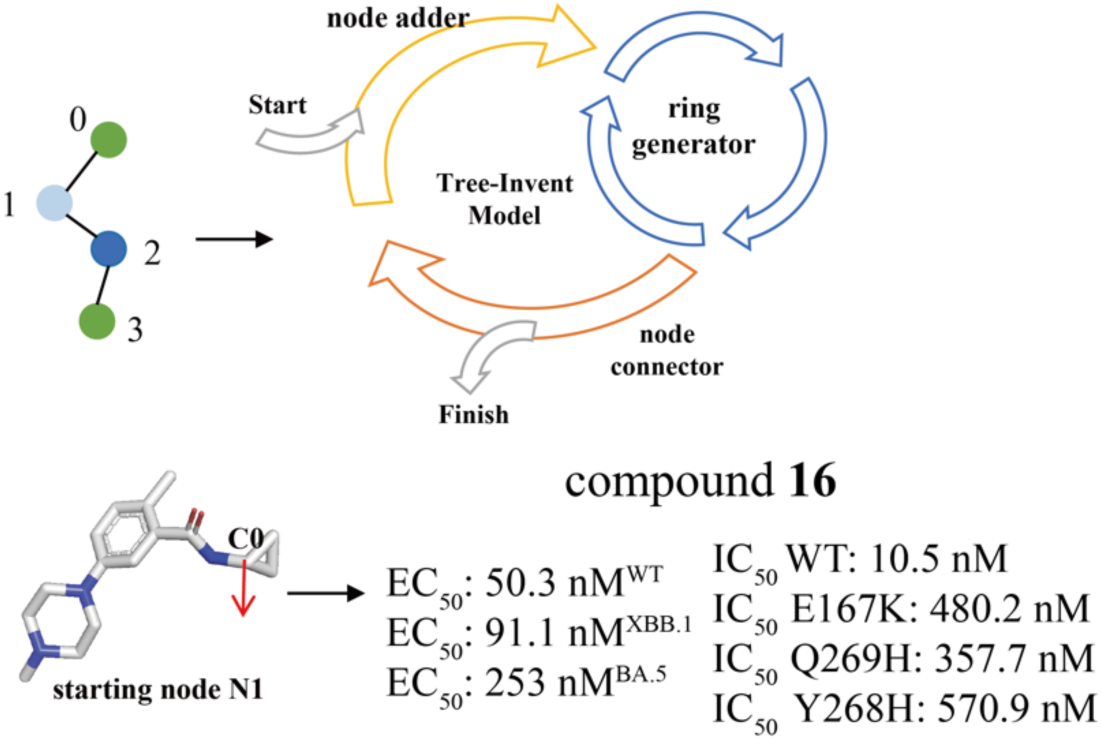

We leveraged an AI generative model to discover a novel PL^pro^ inhibitor with excellent liver stability, low CYP, hERG inhibition and reasonable oral PK properties. At the same time, the compound **16** exhibits high efficacy for the resistance mutation E167K, which resulted in severe resistance to the previously reported inhibitor Jun12682 and PF-07957472.

## Introduction

Severe Acute Respiratory Syndrome Coronavirus 2 (SARS-CoV-2) is the infectious virus responsible for coronavirus disease 2019 (COVID-19). It was initially discovered in China in December 2019 and has rapidly evolved into a global epidemic with a significant cost to human society. Over the past five years, scientists worldwide have been exploring various approaches to treat infected individuals and reduce the severity of the disease. To date, several COVID-19 vaccines have been developed and granted emergency authorization for market distribution^[1]^. All these vaccines target the spike protein to prevent the virus from entering the host cells. However, the high mutation rate of the spike protein, especially within the receptor-binding domain (RBD), diminishes the long-term protective effectiveness of the vaccines^[2]^. This underscores the need to develop alternative therapeutic modalities, particularly small molecule antiviral drugs targeting more conserved viral proteins.

Currently, the FDA has approved three small-molecule drugs, including remdesivir, molnupiravir and nirmatrelvir^[3]^. They all target relatively conserved proteins across different SARS-CoV-2 variants. The first two drugs are viral RNA-dependent RNA polymerase (RdRp) inhibitors, while the third one is an inhibitor of the viral main protease (M^pro^), also known as 3CL protease (3CL^pro^). Remdesivir was the first drug authorized for emergency use, but its administration is limited to intravenous infusion for hospitalized patients with severe symptoms^[4]^. Its clinical efficacy has been found to be sub-optimal, as demonstrated by several clinical trials^[5]^. Similar to remdesivir, molnupiravir is a nucleotide analogue and functions as a mutagen, inhibiting SARS-CoV-2 replication by increasing the viral mutation rate^[6]^. However, there are concerns that molnupiravir may also induce similar mutations in the host^[7]^. Nirmatrelvir, combined with a cytochrome P450 3A (CYP3A) inhibitor (*i.e.,* ritonavir), is an oral medication and suitable for non-hospitalized patients^[8]^. Ensiltrelvir, a M^pro^ inhibitor approved in Japan^[9]^, has demonstrated high efficacy in clinical trials. However, it is also a potent CYP3A inhibitor, which may lead to severe adverse drug-drug interactions with other medications^[10]^. Therefore, new antivirals with alternative mechanisms are still urgently needed.

The papain-like protease (PL^pro^) is one of the two viral proteases responsible for cleaving the polyproteins encoded by the SARS-COV-2 genome^[11]^. Additionally, it can remove the ISG-15 and ubiquitin modifications on host proteins^[12]^. This dual enzymatic function allows it to both promote virus replication and suppress the host’s immune response. Moreover, PL^pro^ is a highly conserved protein across SARS-CoV-2 variants^[13]^, making it a promising drug target for anti-COVID-19 therapy. The catalytic site of the protein is situated within a small buried pocket, adjacent to an induced substrate-binding cavity by the reshaping of the BL2 loop, a flexible beta turn/loop region (Gly266–Gly271)^[14]^.

**GRL0617** was originally discovered as a non-covalent anti-SARS-CoV PL^pro^ inhibitor binding at the induced substrate-binding cavity and later validated to have inhibitory potency against SARS-COV-2 PL^pro^ at the micromolar level^[14]^. Analogs of **GRL0617** were developed with moderate enzymatic and cellular activities, such as **Jun9-84-3**^[15]^, **XR8-23**^[16]^, **Jun11273**^[17]^ and covalent inhibitors **CP7**^[18]^. Recently, a few compounds^[19]^ (such as **Jun12862, PF-07957472, GZNL-P4**) were reported to have in vivo efficacy in SARS-CoV-2 infected mice, along with oral bio-availability. **GZNL-P4** is a potent compound identified in our previous study, but structural optimization is needed to improve its low liver stability. The development of PL^pro^ inhibitors at the clinical stage remains rare. To date, only one PL^pro^ inhibitor has been reported to have entered clinical trials in China, and its structure has yet to be disclosed^[20]^. Therefore, the development of PL^pro^ inhibitors remains necessary to explore their potential as a COVID-19 therapy.

The high mutation frequency of coronaviruses has given rise to a severe and increasingly pressing challenge of drug resistance. During the COVID-19 pandemic, vaccines and antibody drugs played a crucial role; however, later-emerging Omicron subvariants (e.g., BA.1, BA.2, BA.5, BQ.1.1, XBB, XBB.1.5, etc.) exhibited high or complete neutralization escape against nearly all authorized monoclonal antibodies, leading to their successive withdrawal from clinical use ^[21]^. Similarly, mutations in Mpro (e.g., Q192R, H172Y, H172Y/Q189E, L50F/E166V, and L50F/E166A/L167V) have been demonstrated to reduce susceptibility to nirmatrelvir and ritonavir, with the decrease ranging from moderate (25–99-fold) to high (> 100-fold) ^[21–22]^. *In vitro* studies further indicate that multiple mutations in RdRp (including E802D, V166A, N198S, S759A, V792I, C799F/R, etc.) can decrease sensitivity to remdesivir, with the magnitude of reduction varying from several-fold to several hundred-fold ^[23]^. Several key mutations in PL^pro^ have been confirmed to significantly impair the efficacy of existing inhibitors, which constitutes a core challenge for the development of PL^pro^ inhibitors. Tan et al. reported that residues E167, Y268, and Q269 are mutation hotspots associated with resistance to existing PL^pro^ inhibitors ^[24]^. Especially, the mutations of E167K and Y268N in PL^pro^ dramatically decrease the inhibitory activity of Jun2682 and PF-02957472 with the Ki from 26.51 nM and 6.85 nM to greater than 10 μM and 8.036 μM, respectively ^[24]^. Although no PL^pro^ inhibitor has reached clinical approval to date, drug resistance remains a significant concern. Consequently, developing PL^pro^ inhibitors that can overcome drug resistance is critical for future pandemics.

Artificial intelligence (AI) technology has shown great potential to accelerate drug discovery^[25]^. Among these models, deep molecular generation models have garnered particular attention due to their ability to autonomously produce novel molecular structures. Early molecular generation models primarily involved random sampling from the learned continuous molecular chemical space to generate a large number of molecular structures^[26]^. However, due to the vastness of chemical space, the probability of discovering active molecules through sampling these models is relatively low. In recent years, there has been an increasing focus on achieving goal-directed molecular design through the development of conditional molecular generation models. Key approaches include altering the form of model input^[27]^, utilizing conditional deep neural networks ^[28]^, applying transfer learning^[29]^, and employing reinforcement learning^[30]^, etc. Some of the latest methods directly decode the learned three-dimensional structural features of protein pockets into molecular chemical structures^[31]^. Although numerous methods have been developed, only a few of them have been reported with application in real-world scenarios.

In earlier research, we developed Tree-Invent^[32]^, a graph-based molecular generation model with topology constraints. This model is versatile across different drug design scenarios and excels in achieving precise control over structural topology during molecular generation. In this model, a molecular graph is represented as a topological tree comprising ring nodes, non-ring nodes, and edges connecting the nodes. Molecule generation is achieved by carrying out operations of node addition, ring generation, and node connection in an autoregressive manner. Reinforcement learning (RL)^[33]^ is also integrated within the Tree-Invent model to facilitate the exploration of chemical space toward desired properties. We applied this method to conduct scaffold hopping exercise and discovered a novel quinolone chemical series that exhibits nanomolar potency in enzymatic assay and excellent anti-viral potency in cellular experiments. Our best compound exhibits nanomolar level inhibition activity to PL^pro^ E167K and demonstrates reasonable in vitro and in vivo oral PK properties. The significant selectivity over normal human cells and weak inhibition on the CYP and hERG proteins indicate its promising safety profile for antiviral treatments.

## Results

### Scaffold hopping through topology constrained molecular generation with Tree-Invent

So far, exploration of the structure-activity relationship (SAR) based on the structure of **GRL-0617** has resulted in several potent PL^pro^ inhibitors (Fig. S1A). In current study, we utilized the Tree-Invent model (Fig. S2) to design novel PL^pro^ inhibitors, using the crystal structure of PL^pro^ enzymatic domain in complex with known inhibitor GRL0617 (PDB ID: 7JRN), which exhibits potency at micromolar range (Fig. S1B), where the naphthalene ring serves as the head group, is located at the BL2 groove sandwiched between the aromatic residue Y269 and the hydrophobic residues P247 and P248. The aniline group in the tail group is located at the P3 sub-pocket and extends into the P2 sub-pocket near L162 through a methyl group. The linker group is located at the P4 sub-pocket, where the hydrogen bonds with D164 and Q269 are formed by the amide group. Accordingly, the structure of **GRL0617** can be divided into three parts, namely the head, linker and tail part, corresponding to three sub-pockets in the substrate binding site (Fig. S1C) ^[14]^.

In our previous work, we identified a potent compound, **GZNL-P4**, which features a tricyclic dihydroacenaphthylene (DHAN) scaffold. The co-crystal structure of **GZNL-P4** with PL^pro^ (PDB ID: 8YX2) reveals that its tail group extends into the solvent-exposed region at the P3 sub-pocket. The positively charged piperazine nitrogen in this group can form salt bridge with E167, an interaction absent in the **GRL0617** complex structure (Fig. 1A). However, **GZNL-P4** has poor liver metabolic stability (rat liver microsome stability t_1/2_ 7.95 min), which is believed to be associated with the head DHAN group, as observed in the metabolism of **GRL0617** ^[34]^, so finding other head groups is needed to address the stability issue.

**Figure 1.**
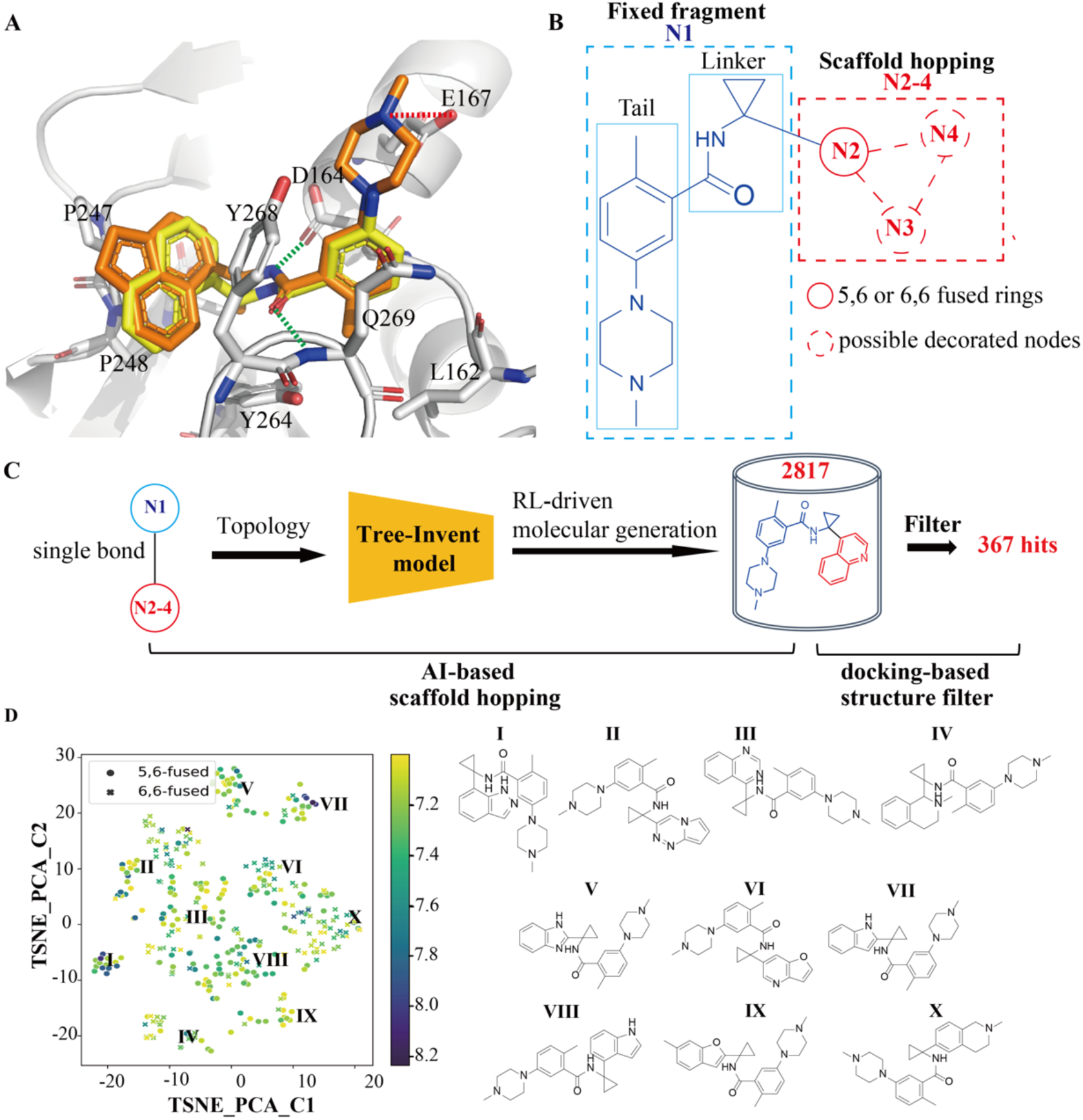
Scaffold hopping of the GZNL-P4 structure. A) Binding mode of **GZNL-P4**. The crystal structures of **GZNL-P4** (orange) and **GRL-0617** (yellow) are overlapped. Ligands and residues surrounding the binding pockets are shown as sticks. Protein secondary structure is shown as transparent cartoons. Green and red dashes represent hydrogen bond and salt bridge interactions. B) Transformation of **GZNL-P4** to a topological tree. The solid and dashed circles represent essential and inessential nodes for molecular generation, respectively. The dashed lines represent the bonds to the inessential nodes. C) Workflow of scaffold hopping of the **GZNL-P4** structure based on the Tree-Invent model using the RL strategy. The blue and red circles represent prefixed and changeable nodes. D) Chemical space analysis based on the PCA analysis ^[37]^ using the Morgan fingerprint ^[38]^. tSNE plot ^[39]^ was used to display the chemical space of molecules filtered based on docking scores (represented by the color of the heatmap) and their pharmacophore model. The labels I-X indicate ten diverse structures with reasonable docking scores.

Herein, we employed Tree-Invent model to explore the chemical space of head group. The structure of **GZNL-P4** was transformed to a topology tree with four nodes connected by single bonds (Fig. 1B). To focus on the generation of the head group, node N1 was predefined as the substructure comprising the linker and tail part of **GZNL-P4** and kept unchanged in the process of molecule generation. Node N2 was designated as a changeable node grown from node A to replace the head group of **GZNL-P4**. Analysis of the binding modes of **GZNL-P4** and **GRL0617** indicates that the aromatic ring system of the head group is important for binding through π-π stacking interaction with Y268 and hydrophobic interaction with P247 and P248 ^[14]^. To mimic this binding mode, node N2 was confined to contain 5,6- or 6,6-fused bicyclic systems with at least one aromatic ring. Furthermore, two nodes, *i.e.,* N3 and N4, were additionally appended to node N2 as R-group decorations. However, their generation was non-essential and entirely contingent upon the model’s decision regarding node addition. With these topological constraints, Tree-Invent was implemented with RL to generate molecules (Fig. 1C), and two rounds of RL were conducted for the generation of 5,6- and 6,6-fused bicyclic molecules. The crystal structure of **GRL0617** was chosen as the protein model for carrying out docking during the generative process and the docking score was transformed into reward score during the RL process, aiming to generate molecules with high affinity.

Through 1000 epochs of RL, the learning curves of the Tree-Invent model converge (Fig. S3A), suggesting the model has learned how to generate high-score molecules with 5,6- and 6,6-fused bicyclic systems. Combining the generation sets of the two RL runs, 2817 unique molecules were obtained. Among them, 1270 molecules have 5,6-fused ring structure and 1547 molecules have 6,6-fused ring structure (Fig. S3B). Moreover, the substructures corresponding to node N1 in these molecules are kept the same as in **GZNL-P4**. Half of the molecules have a naked ring structure in the head group, while the others have no more than two substitutes on the ring structure as expected (Fig. S3C). This demonstrates that the Tree-Invent model is capable of generating molecules satisfying the topological constraints. It was also found that the naphthalene group of **GRL0617** can be reproduced which proves the capability of this method in identifying active molecules. Subsequently, all generated molecules were redocked to the binding pocket using the Glide docking program under the standard precision mode (Glide-SP)^[35]^, resulting to 367 molecules with similar binding mode to **GZNL-P4** and **GRL0617** and docking scores lower than −7.0 kcal/mol, a cutoff to distinguish suitable molecules from all generated molecules (Fig. S3D). Moreover, these remaining molecules are all compliant with the Lipinski rules^[36]^. Chemical space analysis based on the PCA analysis ^[37]^ using the Morgan fingerprint ^[38]^ reveals that these molecules can be grouped into several categories, and each cluster has some molecules with good docking scores (Fig. 1D). The chemical space of molecules filtered based on docking scores was displayed using tSNE plot ^[39]^ (Fig. 1D).

### Identification of novel scaffold of PL^pro^ inhibitors

Considering synthesis feasibility, nine molecules were chosen for synthesis. The synthetic route was illustrated as Scheme 1 in the supplemental text. The structures of the synthesized compounds are shown in Fig. 2A. Their docking poses can be found in Figure S4. Then, they were tested using a fluorescence-based biochemical assay^[40]^ to evaluate their inhibitory activities on PL^pro^ (Fig. 2B). Compounds **1** and **2**, containing a quinoline ring in the head group, are the most active compounds. Their aromatic nitrogen atoms in the quinoline ring are situated at the para-position to the linker substituent. Chlorination on the quinoline ring in compound **1** shows a minor impact on the activity, while compound **2** is suggested to have better binding affinity (Fig. 2C). The binding affinity of compound **2** with PL^pro^ measured by isothermal titration calorimetry (ITC) and biolayer interferometry (BLI) are 285 nM and 187 nM (Fig. S5A, S5C), respectively. Compound **6** with benzothiophene exhibits somewhat lower activity than compound **2**. Other compounds having 5,6 ring systems such as indole, benzopyrazole and 1H-pyrrolopyridine have much worse activity, regardless of whether the head group is a five-membered or six-membered ring. Among them, compound **8** with azaindole is the weakest. N-methyl substitution on the indole ring of compound **9** improves the activity, probably because it makes additional van der Waal interaction with the residue P248. Compound **5** with furopyridine has a special substitution position on the head group, indicating that the direction of substitution on the head group is very important for activity. Through the AI-driven R-group generation process, we successfully identified several PL^pro^ inhibitors with new scaffolds having reasonable potency. The liver metabolic stability of compound **2** was measured, and to our delight, its half-life in liver microsome has been largely improved compared to **GZNL-P4** (Fig. 5E).

**Figure 2.**
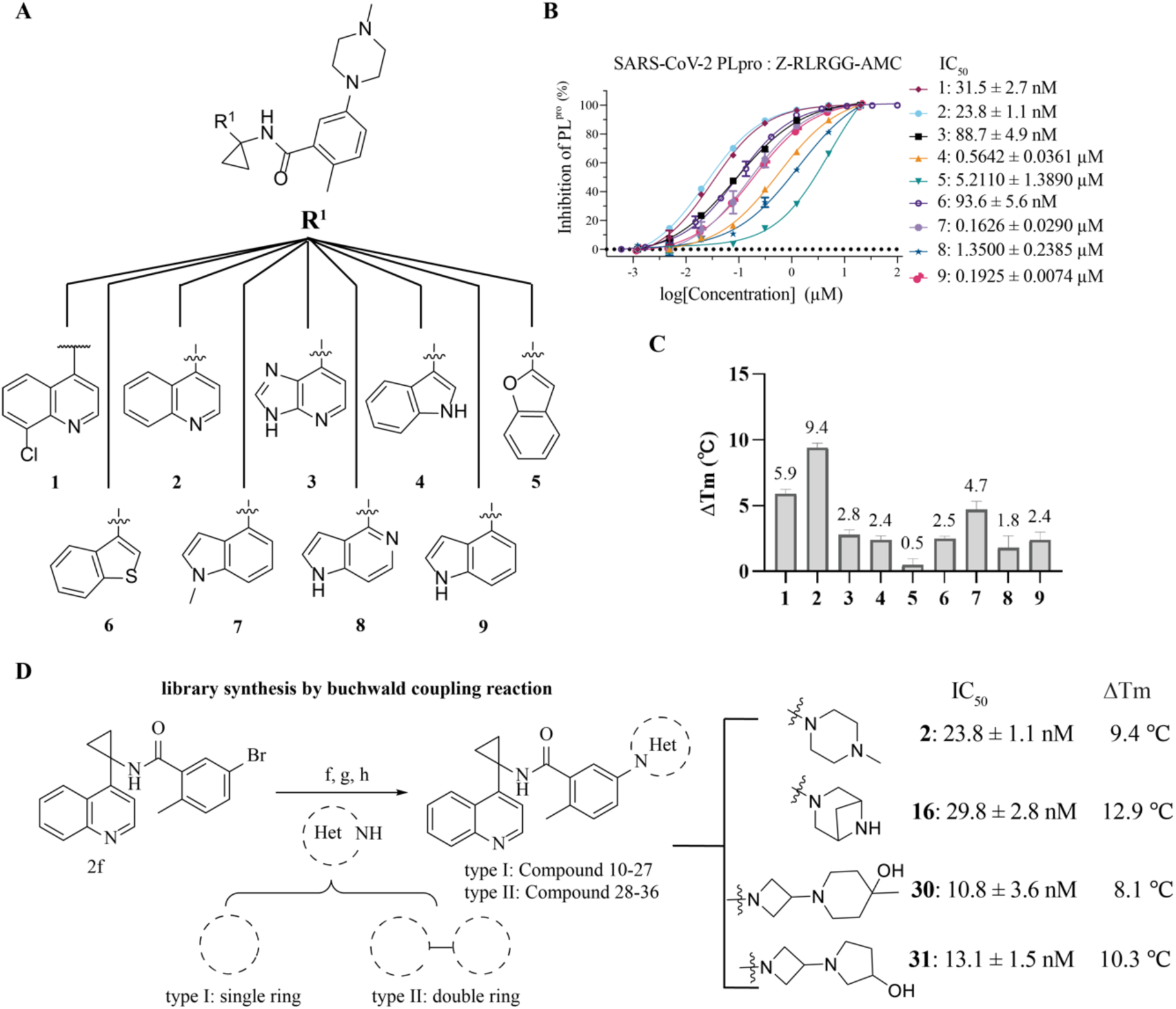
Lead compound identification. A) Structures of nine synthesized compounds **1**-**9**. R^1^ groups are shown at the top. B) Dose-dependent inhibitory activity of compounds **1**-**9** against PL^pro^ as determined by the fluorescence-based biochemical assay. Half maximum inhibitory concentration (IC_50_) is shown in the legend. Data are presented as mean ± s.d. of three technical replicates. C) Differential scanning fluorimetry assay analysis of SARS-CoV-2 PL^pro^ with compounds. The data are presented as the mean of three replicates. D) Scheme of library synthesis for compound **10**-**36**. Synthesis routes were shown in the supplemental text. The inhibition activity (IC_50_) and the delta melt temperature (ΔTm) of lead compounds **2**, **16**, **30**, and **31** were shown. The inhibition activity of IC_50_ is presented as mean ± s.d. of three technical replicates, but the Δ Tm is presented as the mean of three replicates.

To better understand the binding mode of compound **2**, we resolved its co-crystal structure in complex with PL^pro^ (Fig. 3A). The residues surrounding the binding pocket maintain nearly identical conformations to those observed in the co-crystal structure of **GRL0617**, and there is a good alignment between the binding poses of the two compounds (Fig. 3E). The binding mode analysis shows that the quinoline ring of compound **2** is overlapped with the head group of **GRL0617**. As expected, the quinoline ring makes π-π stacking interaction with Y268, and extends the interaction with P247 and P248. The nitrogen atom on the quinoline ring is exposed to solvent, making no obvious binding with protein residues. The cyclopropyl group binds at the bottom of the pocket surrounding Y264 and T301, and the amide linker group makes hydrogen bonds with D164, Y268 and Q269. The tail group, i.e., piperazine-substituted benzyl group, is exposed to solvent, and the positively charged piperazine nitrogen in this group can form a salt bridge with E167, an interaction absent in the GRL0617 complex structure, while the benzyl group extends into the sub-pocket near L162 through a methyl group. This binding mode is consistent with that of **GZNL-P4**, which provides clear guidance for structural optimization.

**Figure 3.**
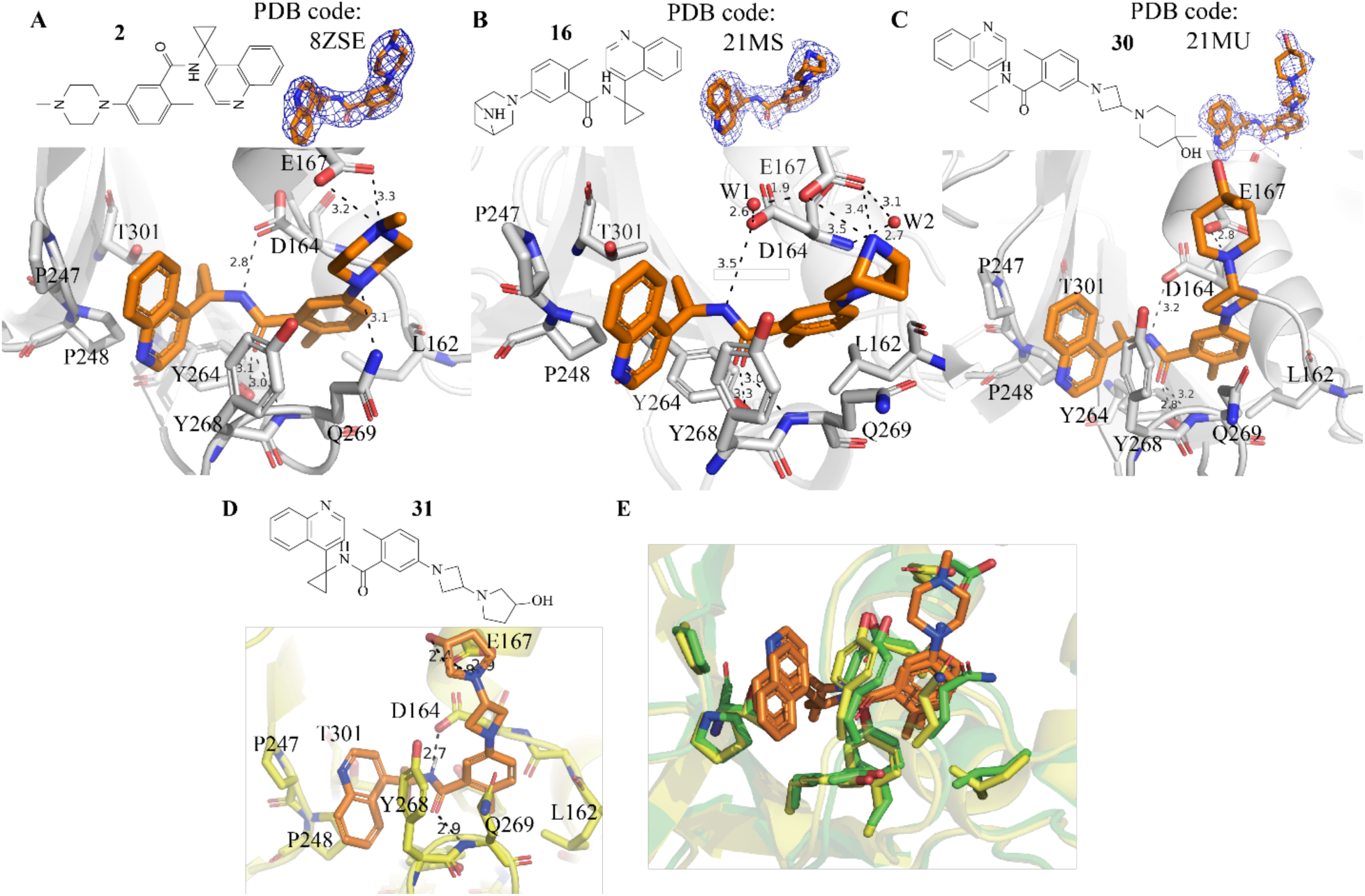
Binding mode of active compounds. A-C) The binding mode of compounds **2** (A)**, 16** (B)**, 30** (C) were determined by X-ray diffraction. Compounds **2, 16, 30** were shown as orange sticks. Residues around the binding pocket are shown as gray sticks. Protein secondary structure is shown as gray cartoons. Black dashes represent hydrogen bonds and salt bridges. The contour level of ligand map was set as 1.0. D) Docking poses of compound **31**. Ligands and residues are shown as orange and yellow sticks, respectively. Black dash represents either hydrogen bond or salt bridge interactions labeled with the distance. E) Alignment of the crystal structures with compound **2** (yellow) and **GRL0617** (green, PDB code: 7JRN). Protein secondary structure is shown as cartoons and residues around the binding pocket are shown as sticks. Ligands are specifically shown as orange sticks.

### Structural optimization of compound 2

To further improve the potency of compound **2**, we explored substituents at the R^2^ position to optimize its interaction with E167. We selected commercially available amine-containing reagents to replace the N-methylpiperazine group of compound **2** and designed a virtual library of 2347 compounds containing positively charged amine groups (Fig. 2D). Molecular docking using the co-crystal structure of compound **2** as docking model was then carried out, compounds whose docking pose is similar to the binding pose of compound **2** and simultaneously maintains the salt bridge with E167 in an appropriate distance (the cutoff is set to 4.0 Å) were selected. At last, 29 top-ranked compounds (by Glide docking score) were selected to synthesize (Scheme 2 in the supplemental text) and their structures are listed in Table 1. They can be divided into two types in terms of the ring count in the R^2^ group.

**Table 1.** SAR exploration of R^2^ group.

| Compound | R<sup>2</sup> | PL<sup>pro</sup> IC<sub>50</sub> (μM) | Compound | R<sup>2</sup> | PL<sup>pro</sup> IC<sub>50</sub> (μM) |
| --- | --- | --- | --- | --- | --- |
| 2 |  | 0.0238±0.0011 | 23 |  | 0.0992±0.0217 |
| 10 |  | 0.0327±0.0016 | 24 |  | 0.0563±0.0059 |
| 11 |  | 0.0288±0.0018 | 25 |  | 0.0156±0.0018 |
| 12 |  | 2.2330±0.2485 | 26 |  | 0.2861±0.0224 |
| 13 |  | 1.0300±0.1295 | 27 |  | 0.3016±0.0287 |
| 14 |  | 0.0453±0.0069 | 28 |  | 0.0376±0.0055 |
| 15 |  | 11.0700±14.6380 | 29 |  | 0.0256±0.0050 |
| 16 |  | 0.0105±0.0006 | 30 |  | 0.0108±0.0036 |
| 17 |  | 0.0316±0.0065 | 31 |  | 0.0131±0.0015 |
| 18 |  | 0.0520±0.0088 | 32 |  | 0.0103±0.0028 |
| 19 |  | 0.0284±0.0022 | 33 |  | 0.0494±0.0019 |
| 20 |  | 0.3836±0.0510 | 34 |  | 0.0519±0.0107 |
| 21 |  | 0.0483±0.0107 | 35 |  | 0.1817±0.0131 |
| 22 |  | 0.0560±0.0090 | 36 |  | 1.0540±0.1536 |

Following SAR analysis reveals that the R^2^ group greatly influences the inhibitory activity of PL^pro^ (Table 1 and Fig. S6). Several compounds show superior activity to compound **2**, and three compounds demonstrate IC_50_ values lower than 0.015 μM. Among type I molecules with a single ring in the R^2^ group, compound **10** with piperazine substitute has worse potency than compound **2** having a methylpiperazine group, which means that the activity is sensitive to the position of methyl substituent on the piperazine ring. Compound **2** with 4-methylpiperazine exhibits much higher activity than compounds **14** and **15** which contain the substitutes of 2- and 3-methylpiperazine. The compounds with hydroxyalkyl groups substituted on the 4-position of piperazine have slightly worse activity than compound **2**. However, substituting with dimethylaminoalkyl groups, as seen in compound **12**, negatively impacts activity. N-sulfonylation of piperazine significantly reduces activity, likely due to the disruption of the salt bridge formed between piperazine and the acid group in E167. It is worthwhile noting that the bridged piperazine significantly enhances activity. Compound **16** (also named as GZNL-2016) is the most potent in this series with IC_50_ of 10.5 nM. Compounds bearing similar cycloamine groups, including the ones with spiral and fused rings, also demonstrate reasonable activity, although they don’t show improvement compared to compound **2**. These groups can mimic the salt bridge interaction of piperazine. Nevertheless, the introduction of seven-member-ring results in a reduction of activity, which may be due to the sub-optimal salt bridge interaction of the secondary amine in the enlarged ring. Compared to compound **2** with a Kd of 285 nM (Fig. S5A), compound **16** exhibits higher binding affinity with PL^pro^. The measured Kd of compound **16** by isothermal titration calorimetry (ITC) and biolayer interferometry (BLI) are 68.6 nM and 78.9 nM (Fig. S5B, S5D), respectively. Type II molecules contain the R^2^ substituents involving two cycloamine groups linked by a single bond. Interestingly, the majority of compounds exhibited comparable activity to compound **2**, except for compounds **35** and **36**. The reduced activity of compound **35** is attributed to the N-methyl-piperazine substituent, which may form a sub-optimal salt bridge with E167. The weak activity observed for compound **36** indicates that piperidine is worse than piperazine for the substitution on the benzene ring, as evidenced by the decrease in the activity of compound **20**. On the other hand, compounds featuring azetidines substituted with pentacyclic/hexacyclic amines demonstrated the highest activity within the sub-group. Notably, compounds **30** (also named as GZNL-2030) and **31** exhibit potency comparable to compound **16**, making them also promising candidates for further profiling. Corresponding to the activity, the binding affinity of compound **30** with PL^pro^ measured by biolayer interferometry (BLI) is 447 nM (Fig. S5E). Most compounds among **10**-**36** can improve the thermal stability of PL^pro^ with a delta Tm larger than 3 ℃ (Fig. S5F). These compounds with larger delta Tm exhibit better potency for PL^pro^ (Fig. S5G). Correspondingly, compound **16** exhibits the largest delta Tm (Fig. S5F), which is consistent with its most potent inhibitory activity against PL^pro^ (Table 1).

The binding poses of the most potent compounds from the crystal structures (compounds **16**, **30**) and docking (compound **31**) are consistent with that of compound **2**, which is characterized by the sandwiched conformation of the quinoline ring, the hydrogen bonds with D164, Y268 and Q269, and the salt-bridge with E167 (Fig. 3A-D). Q269 formed a hydrogen bond with compound **2**, but a π-π stacking interaction with compound **16** (Fig. 3B). In the binding pose of compound **16**, the hydrogen bond network composed of D164, E167 and compound **16** mediated by two water molecules W1 and W2 may lower energy status, which may explain the superior performance of compound **16** to other compounds.

### The enzymatic inhibition of compound 16 on the PL^pro^ mutations

Mutations at residues E167 and Y268 of PL^pro^ lead to a dramatic decrease in sensitivity to the existing PL^pro^ inhibitors Jun12682 and PF-07957472, with a nearly 1000-fold increase in Ki ^[24]^. To assess the susceptibility of compound **16** to distinct PLpro mutations, we evaluated its inhibitory activity against a panel of PLpro mutants, including E167K, E167S, Q269H, Y268N, Y268H, and the double mutant E167G/Q269H (Fig. 4A, S7). The results demonstrated that compound **16** essentially abolished its inhibitory activity against Y268N, while its potency was also significantly diminished against E167S and E167G/Q269H double mutant with IC₅₀ of 4.421 μM and 6.883 μM, respectively. Encouragingly, compound **16** retained relatively high inhibitory activity against E167K, Q269H, and Y268H, with IC_50_ values of 480.2 nM, 357.7 nM, and 570.9 nM, respectively. For compound **16**, the calculated Ki for PL^pro^ E167K using the Cheng-Prusoff equation ^[41]^ was 439.3 nM. In stark contrast, both Jun12682 and PF-07957472 exhibited dramatically declined inhibition activity with Ki around 10 μM against E167K ^[24]^. The binding affinity Kd to PL^pro^ E167K determined using ITC was 256 nM, 1.33 μM, and 1.02 μM, respectively (Fig. 4B-E). This may account for the reason why Compound **16** still retains favorable inhibitory activity against E167K.

**Figure 4.**
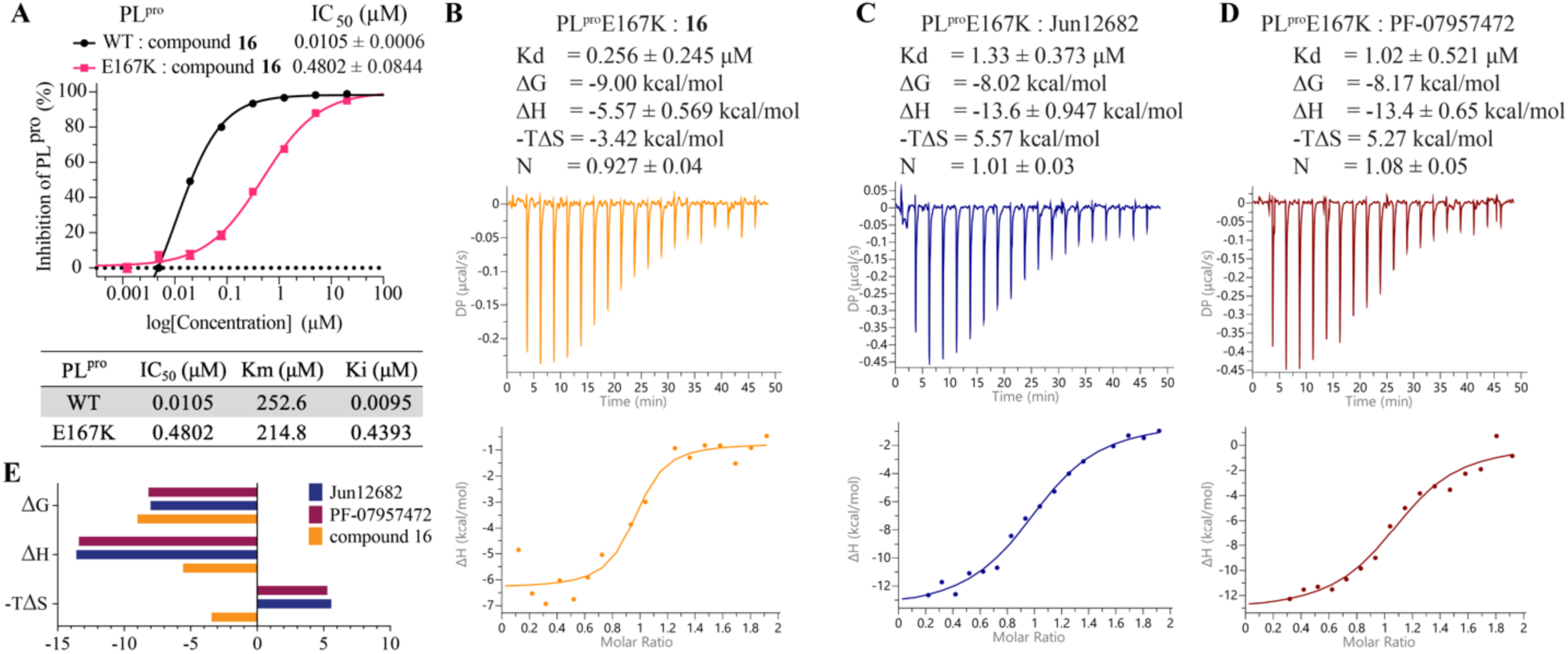
The binding and inhibition activity of compound 16 to PL^pro^E167K mutant. A) The inhibition activity of compound 16 for the PL^pro^E167K mutant conferring drug resistance. The error bars are mean ± s.d based on three technical replicates. B-E) The binding affinity obtained by ITC of compound 16, Jun12682, and PF-07957472 to PL^pro^E167K by ITC.

### Antiviral activity, liver stability and PK profiling of compound 16

The antiviral activities of several potent compounds were measured using a cell-based assay. The antiviral potency with 50% effective concentration (EC_50_) of compound **2** against wild-type SARS-CoV-2 and two epidemic variants Omicron BA.5 and XBB.1 in infected Vero E6 cells are 0.4954 ± 0.1486 μM, 0.6199 ± 0.1061 μM and 0.6159 ± 0.0345 μM (Fig. 5A-C). At the same time, compound **2** shows rather low toxicity to the normal human cell line HEK293T (CC_50_ > 200 μM) (Fig. 5D). Structural optimization on compound **2** resulted in three more potent compounds, compound **16**, **30** and **31**, which have enhanced anti-viral activity and a large enough safety window (CC_50_/EC_50_ for all variants: >379, >292, and >348 folds for compounds **16**, **30**, and **31**, respectively). Notably, compound **16** has the best antiviral activity compared to compounds **30** and **31** on all three tested virus variants (Fig. 5A-C). And, the variant of Omic-BA.5 is the least sensitive to all three PL^pro^ inhibitors.

**Figure 5.**
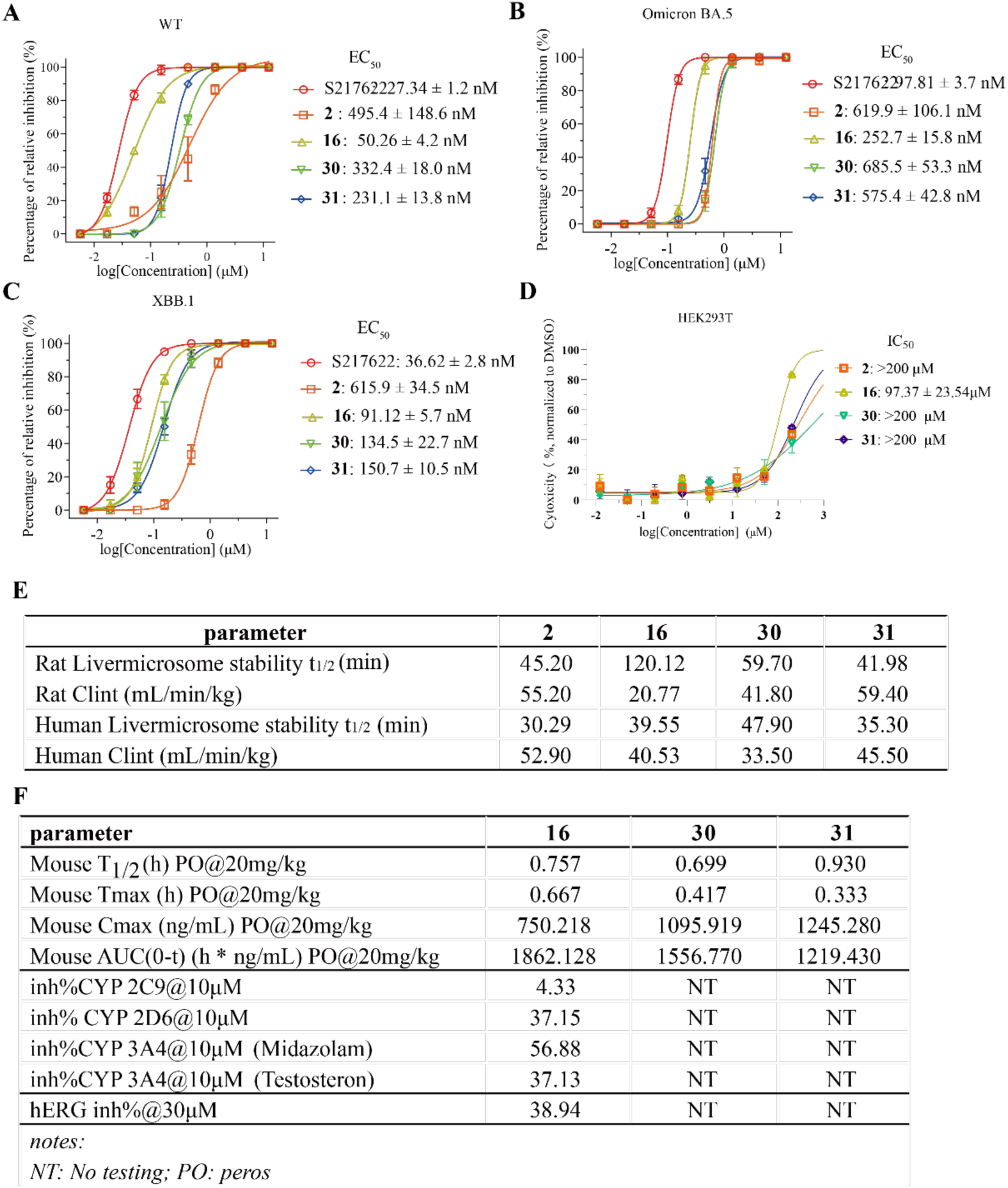
Cellular activity and PK profiling of the compounds 2, 16, 30, and 31. A-C) Antiviral activity of compounds **2**, **16**, **30**, **31** against SARS-CoV-2 wild type (WT) (A), variants Omicron BA.5 (B), and XBB.1 (C). The EC_50_ was measured on virus infected Vero E6 cells. The error bars are mean ± s.d based on three technical replicates. D) Toxicity to normal HEK293T cell lines. Data are presented as the mean of two technical replicates. E) Table for liver stability in rat and human liver microsomes. F) Table for in vivo PK characteristics, CYP inhibition and hERG toxicity.

Compared to the previously reported compound **GZNL-P4** (human and rat liver microsome stability t_1/2_ of 3.8 min and 7.95 min, respectively^[19b]^), the liver stability of the Quinoline series significantly improved the metabolism stability in both rat and human liver microsomes (Fig. 5E). In rat liver microsomal stability assay, compound **16** displayed a prolonged half-life (t₁/₂ = 120.12 min) and a low CLint value (20.77 mL/min/kg), which were 2.66-fold, 2.01-fold, 2.86-fold longer in t₁/₂ and 2.66-fold, 2.01-fold, 2.86-fold lower in CLint than those of compound **2** (t₁/₂ = 45.20 min, CLint = 55.20 mL/min/kg), compound **30** (t₁/₂ = 59.70 min, CLint = 41.80 mL/min/kg), and compound **31** (t₁/₂ = 41.98 min, CLint = 59.40 mL/min/kg), respectively. In human liver microsomal stability assay, compound **16** maintained favorable metabolic performance compared with compounds **2** and **31,** but slightly worse than compound **30**.

*In vivo* PK studies were performed in mice via oral gavage at a single dose of 20 mg/kg, and the key PK parameters were quantified and compared among compounds **16**, **30**, and **31** (Fig. 5F). Compound **16** exhibited significant superiority in systemic exposure, as evidenced by its area under the concentration-time curve from time 0 to the last measurable concentration (AUC₀₋ₜ) was 1862.128 h·ng/mL. This value was 19.6% and 52.8% higher than that of compound **30** (AUC₀₋ₜ = 1556.770 h·ng/mL) and **31** (AUC₀₋ₜ = 1219.430 h·ng/mL), respectively, indicating significantly enhanced oral bioavailability. For the elimination half-life (T₁/₂), compound **16** exhibited a moderate value of 0.757 h, which was comparable to those of compound **30** (T₁/₂ = 0.699 h) and **31** (T₁/₂ = 0.930 h). The time to reach peak plasma concentration (Tmax) of compound **16** was 0.667 h, slightly longer than that of compound **30** (Tmax = 0.417 h) and **31** (Tmax = 0.333 h). However, it still ensured rapid oral absorption and subsequent onset of pharmacological effects. The peak plasma concentration (Cmax) of compound **16** was 750.218 ng/mL, which was lower than that of compound **30** (Cmax = 1095.919 ng/mL) and **31** (Cmax = 1245.280 ng/mL) but still maintained a sufficiently high effective plasma concentration level to elicit the desired therapeutic effects.

In general, compound **16** has the best inhibition activity against the enzyme and virus as well as excellent liver metabolic stability. Based on these favorable properties, compound **16** was selected as the lead compound for further profiling. The CYP and hERG inhibition at 10 μM and 30 μM for compound **16** were summarized in Figure 5F, the results indicate this compound has a low risk of inhibiting selected CYP isoforms and hERG channel. All of these favorable characteristics indicate that compound **16** is a promising lead compound suitable for further optimization.

### *In vivo* antiviral efficacy of compound 16

To assess the *in vivo* anti-viral activity of compound **16**, we treated the model mice infected with SARS-CoV-2 EG.5 by oral administration (Fig. 6A). K18-hACE2 transgenic mice aged 8 weeks were used as our mouse model. Thirty-two female hACE2 transgenic mice were divided into four groups with eight mice in each group to evaluate the efficacy of mock, vehicle, positive control PF-07321332 of 500 milligrams per kilogram (mpk), and compound **16** of 25 mpk in the therapeutic treatment. The weight loss plot shows that the weight loss is less than 5% for all groups (Fig. 6B). The lung live viral titers for the group treated with PF-07321332 at 500 mpk and compound **16** at 25 mpk significantly decreased at 3 days post-infection (Fig. 6C).

**Figure 6.**
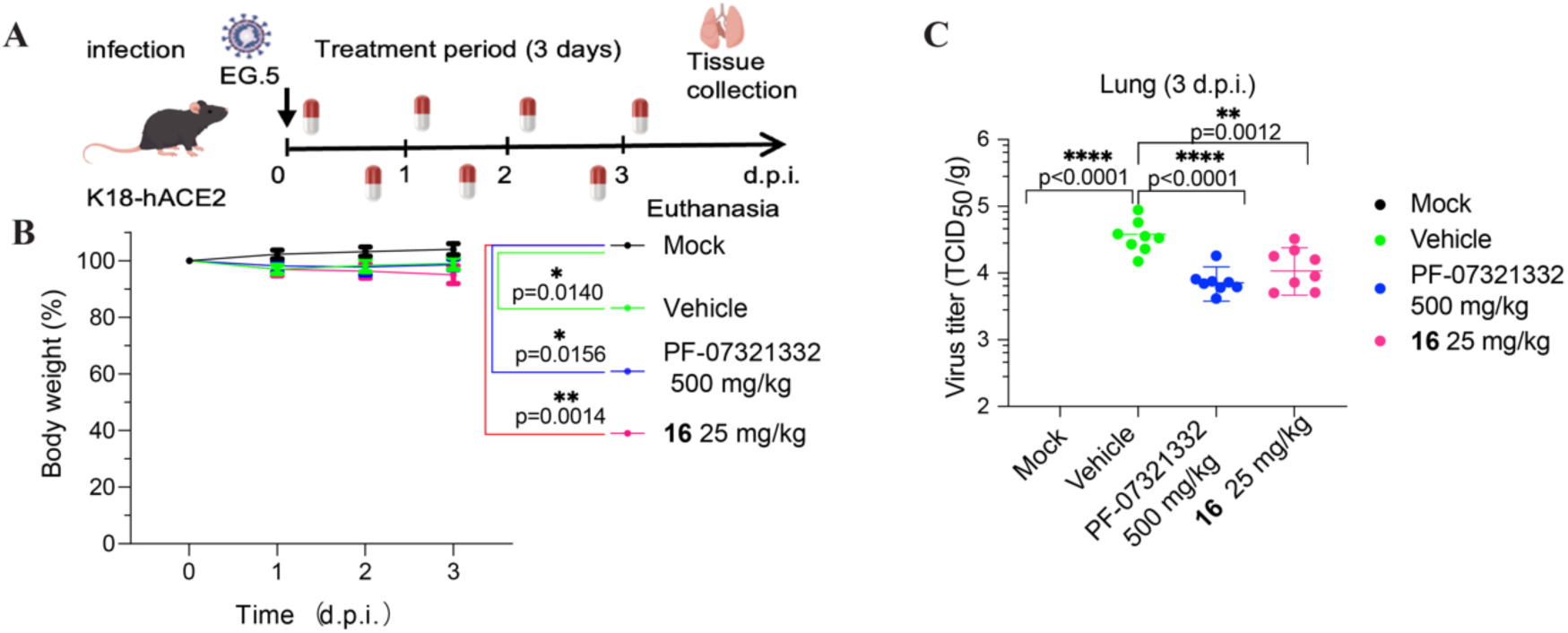
*In vivo* antiviral activity of PL^pro^ inhibitor compound 16. A) Experimental design for the 3-day experiment in K18-ACE2 mouse. B) Body weight loss of mice from different groups (*n* = 8 per group). C) Live viral titers in lungs collected at 3 d.p.i. (*n* = 8 per group). The error bars are mean ± s.d. Statistical differences were determined by Ordinary one-way ANOVA with Dunnett multiple comparisons test for the body weight on day 3 in (B) and the live viral titer in (C). \**P*< 0.05, \*\**P* < 0.01, \*\*\**P* < 0.001, \*\*\*\**P* < 0.0001; ns, not significant.

## Conclusion and Discussion

The continuous mutation of SARS-CoV-2 enables the virus to escape neutralization by antibody therapies^[21]^. Furthermore, resistance to small-molecule drugs is emerging due to mutations in key viral targets such as RdRp, 3CLpro, and PLpro, posing a major challenge to pandemic control. The mutation of 3CL^pro^ at E166V dramatically decreased the susceptibility to Nirmatrelvir (IC_50_ increased 100-fold) ^[42]^. Gandhi et al. reported a De novo emergence of a remdesivir resistance mutation during treatment ^[23a]^. Although no PL^pro^ inhibitors are clinically available, a number of PL^pro^ inhibitors have been reported, such as Jun12682, PF-07957472 ^[19a, 19c]^ and GZNL-P36 ^[19b]^. In addition, drug-resistant mutations of PL^pro^ have also been documented, among which E167 and Y268 are two key drug-resistant mutation sites. In particular, E167G, E167A, E167S, E167K and Y268N confer strong drug resistance to Jun12682 and PF-07957472. Specifically, the Ki value of PF-07957472 increases by more than 130-fold against E167K and 969-fold against Y268N; the Ki value of Jun12682 increases by more than 130-fold against E167K and 385-fold against Y268N. Thus, the drug development targeting the E167 and Y268 mutations of PL^pro^ will help address the potential drug resistance that may arise in the future.

In the current study, we leveraged an AI generative model to discover a novel series of PL^pro^ inhibitors targeting the SARS-CoV-2 virus. Using the generative model, Tree-Invent, we successfully identified a novel scaffold of PL^pro^ inhibitors exhibiting comparable activity to a known reference compound **GZNL-P4**, though it has much improved liver metabolic stability. Further optimization work on this series results in a lead compound **16** with an improved enzyme potency and cellular-level antiviral activity. This compound demonstrates excellent liver stability, low CYP and hERG inhibition and it also demonstrates reasonable oral PK properties. At the same time, unlike the previously reported PL^pro^ inhibitors Jun12682 and PF-07957472 ^[24]^, compound **16** exhibits strong activity against E167K and can effectively overcome the drug resistance caused by the E167K mutation. We believe that this study offers valuable insights for the future development of PL^pro^ inhibitors as a potential treatment for COVID-19 disease, especially provides a scaffold compound and points the way forward for overcoming E167K-mediated resistance.

## Methods and Materials

Detailed descriptions of the in vitro pharmacology studies, *in vivo* pharmacology studies, transcriptomics studies, X-ray crystallography, computational study, and synthetic methods can be found in Supplementary Information.

## Data analysis

The data in the figures represent mean ± SD. All data were analyzed using GraphPad Prism 9.0 software. Statistical comparison between different groups was performed using the corresponding statistical analysis labeled in the figure legends combining several experiments. The binding mode of ligands with protein was illustrated by PyMOL2.6.0a0 (open source).

## Data Availability

Crystal structures generated during the current study are available in the Protein Data Bank (PDB) under accession codes 8ZSE (PL^pro^ bound to compound **2**, here renamed as GZNL-2002), 21MS (PL^pro^ bound to compound **16**, here renamed as GZNL-2016), and 21MU (PL^pro^ bound to compound **30**, here renamed as GZNL-2030). The supplementary note for synthetic compounds can be found in Supplementary Information.

## Acknowledgments

This work is supported by the fundings: Overseas Experts Supporting Programs under National Research Platform (WGZJ22-001); National Natural Science Foundation of China (82170473); The Major Program of Guangzhou National Laboratory (GZNL2024A01005, GZNL2023A01008); Guangdong Natural Science Foundation (2021QN020451, 2021CX020227, 2024A1515011589); Basic and Applied Basic Research Projects of Guangzhou Basic Research Program (2023A04J0161). We thank the staff at Guangzhou Laboratory Core Facility and Animal Center. We also thank the staff at BL10U2/BL02U1/BL17B1/BL18U1/BL19U1 beamlines at Shanghai Synchrotron Radiation Facility (SSRF) of the National Facility for Protein Science in Shanghai (NFPS), Shanghai Advanced Research Institute, Chinese Academy of Sciences, for providing technical support in X-ray diffraction data collection and analysis.

## Author Contributions

J.S, Y.L., Q.Y., W.L., M.D., J.T., J.C., J.Z., K.W., and X.W. performed the cellular and biochemical assays. Y.L. and J.S. performed crystallography. G.Z., P.Z., H.C., P.H., and T.R. performed the medicinal chemistry. T.R., M.X., M.T., and C.H. performed the computational studies. Y.L., W.L., M.D., J.C., J.Z., K.W., and X.W. performed cloning, protein purification and biophysical analysis. Y.L., Q.Y., T.R., G.Z., X.C., C.H., and J.S. conceived the experiments and wrote the manuscript with input from all authors.

