## Supporting Information for "Discovery of novel quinoline papain-like protease inhibitors for COVID-19 through topology constrained molecular generative model"

### Materials and Methods

Our research complies with all relevant ethical regulations. Animal Ethics Committee at Guangzhou National Laboratory and Pharmaron's Institutional Animal Care and Use Committee (IACUC) that approved the study protocol.

### General methods for chemical synthesis

Synthesis routes: compound preparation and structure characterization were elaborately stated in supplemental information. All compounds were isolated through column chromatography or preparative liquid chromatography equipped with a PDA detector and a SB-C18 column (30 × 150 mm) of waters. The mobile phase consisting of buffer A (ultrapure $H_2O$ containing 0.1% formic acid) and buffer B (chromatographic grade $CH_3CN$) was applied with a flow rate of 20 mL/min. All solvents and chemicals were used without further purification. $^1H$ and $^{13}C$ NMR spectra were recorded on a Bruker Avance neo at 600 MHz for $^1H$ NMR and 100 MHZ for $^{13}C$ NMR, respectively. Chemical shifts (δ) were expressed in parts per million (ppm) with TMS as an internal standard. The letters s, d, t, and q represent multiplet. The purity of all tested compounds was larger than 95%, as estimated using a Waters Acquity UPLC system equipped with a PDA detector and a BEH-C18 column (2.1 × 50 mm). The mobile phase consists of buffer A (ultrapure $H_2O$ containing 0.1% formic acid) and buffer B (chromatographic grade $CH_3CN$) and the flow rate is 0.4 mL/min.

### Molecular generation with the Tree-Invent model

Tree-Invent is a topology constrained molecular generative model recently developed by our group[32]. The unique feature of the algorithm is utilizing a topological tree structure as a constraint for molecular generation (Figure6A). It is particularly suitable for lead optimization tasks, such as scaffold hopping, R-group decoration etc, when only part of structure needs to be changed. Tree-Invent integrates the topological tree into an atom-based autoregressive structural generation process (For more details, please refer to [32]), where each node of the topological tree is sequentially decoded to either a single atom or a ring structure. Compared to available models specific for lead optimization, such as SAMOA[38], GraphGMVAE [39], and DeepHop [40], Tree-Invent provides a direct control on molecular generation.

The architecture of Tree-Invent contains three independent modules to complete molecular generation, namely node adder, ring generator and node connector (Figure6A). The module of node adder determines the type of a node and whether the structure should be terminated at current status. The module of ring generator is executed only when a ring node is sampled. This module provides precise control on the generation of ring structure through constraining ring size, number of ring, and number of aromatic ring in the ring system. The module of node connector determines the bond type between the existing node and the new node and also the atom position in the existing node for node growth or connection. Particularly, the model can generate structures according to given topological constraints through masking the action distribution probability (ADP) [32]. Through this unique masking mechanism, Tree-Invent can not only imposes precise control on generation of certain substructures but also allows generation of other parts of the structure in an uncontrolled manner. Furthermore, by incorporating reinforcement learning, Tree-Invent can achieve goal-directed structure generation by optimizing the reward for a given structure.

In this study, we transformed the 2D structure of a reference compound **GZNL-P4** into a simple topological tree (as shown in Figure3B). One pre-defined node (i.e. node N1) represents the structure comprising the linker and tail parts of **GZNL-P4**. Among the atoms in the node, the atom C0 is regarded as the connection atom with new node N2 (Figure6B), while other atoms are blocked to prevent connecting the new node by including them in the predefined saturation atom list (for configurating saturation atom list in node definition, please refer to supplemental material), which the connecting probability is masked to zero. These settings enable node N1 to be completely fixed in the process of molecular generation. Other parts of molecule including node N2, N3 and N4 are specified to only grow from node N1 sequentially, starting with a single bond, mimicking the head group of reference **GZNL-P4**. More specifically, node N2 is confined to have bicyclic ring systems, which contains at least one aromatic ring and at most two R-group substitutions. In the auto-regressive process of molecular generation, maximal node adding operation was set to four steps, in which the first two steps were specified to cover the pre-defined node N1 and bicyclic node N2. In the two additional steps (N3 and N4 node), adding substitutions on the bicyclic ring is done in uncontrolled mode (ie. no masking on the ADP) and their generation is purely dependent on the current state of the structure (Figure6B), which means that the generation of these two nodes may be skipped by the model to generate naked ring as the head group. These topology constrains were pre-defined in the job configuration file.

In total, 1000 epochs of RL were run to generate molecules by rewarding molecules for better docking score. The prior model for RL was trained on the GuacaMol dataset [32] and in the process of RL, the learning rate was set to 0.00005 with the batch size of 20. The range of raw docking score from -8 kcal/mol to -5 kcal/mol was used to define the Sigmoid transformation function of docking score P ($P \in [0, 1]$), which is included in the scoring function of RL. DockStream[41], a docking wrapper to facilitate *de novo* molecule design, was integrated in the Tree-Invent model to integrate Glide docking using the high-throughput mode (Glide-HTVS)[35].

**Molecular docking**

Molecular docking was conducted utilizing the Glide docking program of Schrödinger software suite (version 2020)[27]. The co-crystallized structure of the SARS-CoV-2 PL$^{pro}$ enzyme in complex with **GRL0617** (PDB code: 7JRN) was served as the protein model for the docking. The protein structure underwent pre-processing using the Protein Preparation Wizard tool within the software, ensuring proper assignment of bond orders and repair of amino acids with missing side chains. Additionally, all crystal waters were removed. Subsequently, the Epik algorithm[42] was employed to generate ionization and tautomeric states of the protein at a pH of 7.0. The structure was then minimized using the OPLS3 force field[43] until heavy atoms converged to a root mean square deviation of 0.3 Å from the original coordinates. The minimized structure was utilized to prepare the receptor grid for Glide docking. Concurrently, ligands intended for docking were prepared using the LigPrep module in Schrödinger. Analogous to protein preprocessing, ligands were protonated using the Epik algorithm, and tautomeric and stereoisomeric forms were enumerated. For each compound, a single low-energy conformation was generated utilizing the OPLS3 force field. The prepared ligands were then docked into the predefined docking site within a box size of 20 Å. To evaluate the generated molecules, the docking process employed standard precision, allowing for flexible ligand conformation sampling. After docking, a maximal 10 poses were retained for every ligand and ranked based on the Glidescore. Poses with a docking score lower than -7.0 kcal/mol were retained. Besides, poses that did not fit to the binding mode of **GRL0617** or compound **2**, particularly the stacking interaction with Y268 and P248/P247, as well as hydrogen bonding with D164 and Q269 mediated by the amide group, were eliminated.

**Plasmid Construction, Protein Expression and Purification**

The papain-like protease domain sequence was retrieved from the complete genome of SARS-CoV-2, accessible in the NCBI gene databank under the Gene ID: 43730578. Subsequently, the protein sequence encoding the Ubl domain of the Nsp3 protein (spanning amino acids 746-1060, accession number YP_009742610.1) was codon-optimized for enhanced expression in bacterial systems. The optimized gene was then chemically synthesized and cloned into the pET28a expression vector with BamHI and XhoI (Tsingke). A recognition sequence for the TEV protease was strategically inserted between the N-terminal His6-tag and the $PL^{pro}$ sequence to facilitate subsequent tag removal.

The $PL^{pro}$ mutants C111S ($PL^{pro}$C111S), E167K ($PL^{pro}$ E167K), E167S ($PL^{pro}$ E167S), Q269H ($PL^{pro}$ Q269H), Y268N ($PL^{pro}$ Y268N), Y268H ($PL^{pro}$ Y268H) and E167G/Q269H ($PL^{pro}$ E167G/Q269H) were generated by site-directed mutagenesis using PCR. For the protein expression, BL21(DE3) *Escherichia coli competent cells* were transformed with the $PL^{pro}$ expression plasmid. These cells were then inoculated into LB medium and grown to an OD600 of 0.6-0.8. Subsequently, protein production was induced by adding 0.5 mM of isopropyl-D-thiogalactopyranoside (IPTG) and 1mM zinc chloride ($ZnCl_2$) at 16°C for 18 hours. After induction, the cell pellets were collected through centrifugation at 4000 rpm for 30 minutes and stored at −80 °C for future use. For protein purification, the frozen cell pellets were thawed and resuspended in buffer A, containing 50 mM Tris-HCl, 150 mM NaCl, 10 mM imidazole, and 1 mM TCEP (pH 7.4), supplemented with lysozyme, deoxyribonuclease, and a protease inhibitor cocktail (Roche) to protect the target protein from degradation. Cell lysis was achieved through sonication, and the lysates were separated from cellular debris by centrifugation at 18,000 × g for 35 minutes, repeated twice at 4 °C. The clarified supernatant was further filtered using a 0.22 μm pore size membrane to remove any remaining particulate matter. Subsequently, the supernatant containing His-tagged target protein was loaded onto a HisTrap HP column (5 ml; GE) pre-equilibrated with buffer A. After removal of nonspecific binding proteins by washing with buffer A, the target protein was eluted with a linear gradient of buffer B, containing 50 mM Tris-HCl, 150 mM NaCl, 250 mM imidazole, and 1 mM TCEP (pH 7.4) over 20 column volumes.

To remove the His-tag, the eluted protein was dialyzed in the presence of TEV protease. Following the removal of His-tag, the protein solution was concentrated to 2 ml and loaded onto a Superdex 75 column (120 ml; GE) that had been pre-equilibrated with buffer C, containing 20 mM

Tris-HCl, 100 mM NaCl, and 1 mM TCEP (pH 7.4). This final chromatographic step was performed to achieve high purity and homogeneity of the target protein, ready for downstream applications.

**PL$^{pro}$ enzymatic assay**

The activity of PL$^{pro}$ was assessed using the labeled peptide Z-RLRGG-AMC (GLPBIO, GA23715) as a substrate in a 384-well plate (GREINER, 784076), with the excitation and emission wavelengths at 340 nm and 460 nm. The enzymatic reaction was carried out in the buffer containing 50mM HEPES, 10mM DTT, 0.1mM EDTA, 0.005% tween 20, pH7.2, with GRL0617 (TargetMol, Shanghai, China) as the positive control. The serially diluted compounds were added into PL$^{pro}$ or PL$^{pro}$ mutations solution diluted with assay buffer and incubated for 10 minutes. Then, the diluted substrate peptide with assay buffer was added into PL$^{pro}$ and inhibitor mix solution and incubated for 1 hour at 37 °C. The final concentrations of components in the reaction system are 10 nM and 20 μM for PL$^{pro}$ and substrate, respectively. Fluorescence intensity was recorded with a BioTek Neo2 multimode plate reader (Agilent). The inhibition activity $IC_{50}$ values were calculated using nonlinear regression fitting in GraphPad Prism 9.0. The Ki of compound **16** to PL$^{pro}$ E167K was calculated using the Cheng-Prusoff equation from the value of $IC_{50}$.

**Differential scanning fluorimetry assay (DSF)**

The fluorescence dye SYPRO Orange (Sigam-Aldrich, S5692) was used as a fluorescence probe in a differential scanning fluorimetry assay. The protein sample was diluted to 10μM in PBS (pH7.4) and incubated with 20 μM of compounds, with DMSO as a control. The fluorescence dye SYPRO Orange (Sigam-Aldrich, S5692) was then added to the protein samples at a 5x dye concentration. The sample was mixed thoroughly and transferred to the 384-well microplate (10 μL per well). Protein buffer containing DMSO was used as a blank control. The microplate was placed in the fluorescence quantitative PCR instrument. Fluorescence was monitored when the temperature was gradually raised from 30 to 80 °C in 0.2 °C increments at 5-second intervals. Three technical replicates were tested for each sample. Melt curve data were plotted using the Boltzmann model to obtain the melt temperature (Tm, the midpoint of unfolding of the protein) using GraphPad Prism 9.0.

**Bio-layer interferometry (BLI)**

The BLI assay was performed at 25 °C using the Octet R8 instrument (Sartorius) with phosphate buffer solution (PBS) supplemented with 0.005% Tween-20 (PBST). Biotinylated SARS-CoV-2 PL$^{pro}$ protein was prepared using a commercial biotinylation kit (G-MM-IGT, Genemore). The PL$^{pro}$ protein was immobilized onto Super Streptavidin SSA capture biosensors (18-5070, Sartorius), which were pre-equilibrated with PBST. Subsequently, the biosensors were exposed to varying concentrations of the test compound solution for the association, followed by dissociation in PBST. Data acquisition and analysis were conducted using the Octet® BLI Analysis software.

**Isothermal titration calorimetry (ITC)**

The ITC experiment was carried out at 25 °C using a MicroCal PEAQ-ITC instrument (Malvern Panalytical). The titration protocol consisted of 19 injections, including an initial pre-injection of 0.4 μL followed by 18 injections of 2 μL. The reference power was set to 5 μcal/s, and the compound solution was titrated at intervals of 150 s. Both the $PL^{pro}$ protein and compound were diluted with a buffer (*p*H 7.4) composed of 20 mM Tris-HCl, 100 mM NaCl, 1 mM TECP, and 0.5% (v/v) DMSO. Prior to titration, all samples were centrifugated at 18000 × g for 10 minutes to remove precipitation and air bubbles. The $PL^{pro}$ protein was injected into the sample cell, while the compound was placed into the syringe cell. The reference cell was filled with deionized water as a heat balance control. The collected ata were fitted into a one-site model, and the thermodynamic parameters, including $K_d$, N, ΔG, ΔH, and −TΔS values were calculated using MicroCal PEAQ-ITC analysis software.

**Crystallization, data collection and structure determination**

The purified SARS-CoV-2 $PL^{pro}$C111S protein was incubated overnight with compounds **2, 16 and 30** at a 1:3 protein-to-ligand molar ratio in buffer C, and concentrated to 8 mg/ml. Crystal screening was carried out using the sitting-drop vapor diffusion technique at 4 ℃ in 96-well crystallization plates. Each crystallization drop contained a 1:1 mixture of the protein complex sample and reservoir solution. The protein-ligand complex crystals were obtained after 7-10 days after the droplet setting. Optimal crystallization conditions were determined: 0.05 M Zinc acetate dihydrate, 22% w/v Polyethylene glycol 3,350 for compound **2;** 0.2 M MgCl2.6H2O, 0.1 M BIS-TRIS pH 6.5, 25% w/v PEG 3,350 for compounds **16 and 30**. X-ray diffraction data were collected at the Shanghai Synchrotron Radiation Facility's beamlines BL02U1. Data processing was performed using the Aquarium program [44]. The protein structures were solved by molecular replacement using the program Phaser [45] integrated in PHENIX package [46], employing the previously published $PL^{pro}$ structure (PDB code: 7CJD and 8YX5 for **2** and **16**/**30,** respectively)[47] as a search model. Structure refinement was carried out in PHENIX, with manual interactive model rebuilding conducted in Coot [48]. Ligand constraints were generated using Phenix.elbow [49] or CCP4.AceDRG {Long, 2017 #62}, and ligand models were built based on the omit map. The data collection and refinement statistics are summarized in Table S1.

**Cell culture and cell viability assay**

African green monkey kidney cell Vero E6 (ATCC, CRL-1586) and human embryonic kidney cell HEK293T (ATCC CRL-3216) cells were maintained in Dulbecco's Modified Eagle Medium (DMEM) enriched with 10% fetal bovine serum (FBS), 100 IU/mL penicillin, and 100 μg/mL streptomycin. Cells were incubated at 37 °C in a humidified atmosphere of 5% $CO_2$ and confirmed to be free of mycoplasma contamination. Cell viability was assessed using the Cell Titer-Glo 2.0 Cell Viability Assay (G9242, Promega) following the manufacturer's protocol. Briefly, $4 \times 10^3$ cells suspended in 20 μL of culture medium were plated in opaque-walled 384-well plates and allowed to adhere for 12 hours. After a 48-hour treatment with compounds, 20 μL of Cell Titer-Glo reagent was dispensed into each well. Following 2 minutes of shaking and 10 minutes of incubation, luminescence was quantified using a BioTek Synergy H1 multimode reader (Agilent).

#### Virus preparation and titrations

SARS-CoV-2 wild-type strain (WT, GenBank: MT123291), the Omicron BA.5 variant (Omicron BA.5, GDPCC-303), the XBB.1 variant (XBB.1, IQTC-1596943) and EG.5 variant (IQTC-IM23676) were preserved in Guangzhou Customs Inspection and Quarantine Technology Center (IQTC) BSL-3 Laboratory. The above-mentioned virus strains were propagated in Vero E6 cells using established protocols [50] and stored at -80 °C for future use. Virus titers were determined by performing 10-fold serial dilutions in confluent Vero E6 cells seeded in 96-well microtitre plates. After three days of post-inoculation, a cytopathic effect (CPE) was observed and scored, and the Reed-Muench formula was employed to calculate the TCID50, representing the tissue culture infectious dose causing CPE in 50% of the cell cultures. The infection experiments were conducted within a Biosafety Level 3 (BSL-3) facility at the Guangzhou Customs Inspection and Quarantine Technology Center (IQTC), adhering to strict safety protocols to ensure the safety of personnel and the environment.

#### In vitro antiviral activity assay

The in vitro antiviral activity was performed in Vero E6 cell lines using the CPE method [50]. A 96-well plate was seeded with $2 \times 10^4$ Vero E6 cells and incubated for 24 hours to allow for cell attachment and monolayer formation. Subsequently, various dilutions of the compounds were combined with SARS-CoV-2 at a multiplicity of infection (MOI) of 0.01. Each well received 200 μL of this compound-virus mixture, which was inoculated onto the monolayer of Vero E6 cells. Seventy-two hours post-inoculation, the CPE was quantified using a Celigo Image Cytometer. The inhibitory activity of the compounds was determined based on the CPE rates observed for SARS-CoV-2. The $EC_{50}$ value, representing the effective concentration at which 50% of the CPE is inhibited, was calculated from these data. Each experiment was conducted independently three times, with eight concentration gradients tested in triplicate wells. Representative data from one of these experiments are presented.

#### SARS-CoV-2 infection in K18-hACE2 mice

K18-hACE2 transgenic mice aged 8 weeks were obtained from GemPharmatech. The use of K18-hACE2 transgenic mice has received ethical approval from the Animal Ethics Committee at Guangzhou National Laboratory (GZLAB-AUCP-2025-07-A01). The experiments of SARS-CoV-2 infection in K18-hACE2 mice were conducted within a Biosafety Level 3 (BSL-3) facility at the Guangzhou Customs Inspection and Quarantine Technology Center (IQTC), adhering to strict safety protocols to ensure the safety of personnel and the environment. Thirty-two female hACE2 transgenic mice were divided into four groups with eight mice in each group to evaluate the efficacy of compound **16** (also named GZNL-2016) in the therapeutic treatment. On the day of infection, the hACE2 mice were intranasally inoculated with either $8.9\times10^3$ $TCID_{50}$ EG.5, pre-diluted in 50 μL DMEM. Treatment was delayed until 2 hours post-infection (h.p.i.). K18-hACE2 transgenic mice were orally administered a dose of 500 mg/kg PF-07321332 (InvivoChem, V2402-1g) and 25 mg/kg **16.** The drugs were diluted in 200 μL 5% DMSO/20% hydroxypropyl-betacyclodextrin for the treatment group or vehicle solution only for the control group. Mice were killed at the designated time points and organ tissues were sampled for virological analysis.

**Metabolic stability in liver microsome**

To assess the in vitro metabolic stability, a series of incubations were conducted using rat and human liver microsomes. Each compound was diluted to a concentration of 1.5 µM and incubated with either rat or human liver microsomes (0.75 mg/mL) in a 0.5 mL volume of PBS. Assay plates were prepared with 30 µL aliquots of the compound-microsome mixture, designated for various time points (0, 5, 15, 30, and 45 minutes). The reaction was initiated by adding NADPH to a final concentration of 2 mM, and the mixtures were incubated at 37 °C. At each designated time point, the reaction was terminated by adding an acetonitrile solution containing internal standard (IS, Tolbuamide). The plates were then shaken at 600 rpm for 10 minutes followed by centrifugation at 6000 rpm for 15 minutes. This centrifugation step separated the precipitated protein from the supernatant, which was then transferred to a 96-well plate containing 140 µL of pure water. The samples were analyzed using LC/MS (Shimadzu Nexera LC-40 & SCIEX TQ-6500+) to quantify the remaining concentrations of the compounds. The intrinsic clearance, representing the rate of substrate depletion, was calculated by considering the amount of microsomes per liver weight and the liver weight per kilogram of body weight.

**In vivo PK study**

In this study, pharmacokinetic assessments were conducted on SPF male ICR mice (n = 3) at Medicilon (Shanghai, China). The PK studies were approved by Medicilon's Institutional Animal Care and Use Committee (IACUC) with the ethical approval number 19250-23024-NG. The mice were dosed with 20mg/kg of the test compound using an oral administration route. The dosing vehicle comprised a solution containing 5% DMSO, 10% of the test compound, and 85% saline. Sequential blood samples were collected from the mice at various time points post-administration: 0.25h, 0.5h, 1h, 2h, 4h, 6h, 8h, and 24h. These samples were then analyzed using liquid chromatography tandem mass spectrometry (LC-MS/MS) to quantify the concentration of the compound in the blood. To assess the pharmacokinetic behavior of the compound, key parameters were calculated using Phoenix WinNonlin 7.0 software. These parameters included the area under the concentration-time curve from time zero to the last measurable concentration ($AUC_{0\text{-}t}$), the maximum plasma concentration ($C_{max}$), the time to reach Cmax ($T_{max}$), and the elimination half-life ($T_{1/2}$).

**CYP inhibition**

The compound was added into the liver microsomes solution pre-warmed at 37 °C for 5 min. Subsequently, 20 µL of a pre-warmed (37 °C) 10 mM NADPH solution was introduced into 180 µL of the liver microsomes solution containing the compounds, thus initiating the enzyme reaction. The assay plate was then incubated at 37 °C until the designated time point, at which 300µL of a termination solution containing 35 ng/mL of ketoprofen, 7.5 ng/mL of carbamazepine, 5 ng/mL of diphenhydramine, and 10 ng/mL of tolbutamide was added to halt the reaction. Following termination, the samples were centrifuged at 3,220 × g for 40 minutes. The supernatant was carefully transferred to an analysis plate containing an appropriate volume of ultra-pure water, ready for LC-MS/MS analysis. Through this process, the inhibition rate was determined for the CYP isoforms 2C9, 2D6, and 3A4.

**hERG inhibition**

A patch clamp system procured from Sutter Instrument (USA) was employed to monitor the current in human embryonic kidney cells stably expressing hERG channels (hERG-HEK293 cells, Creacell, A-0320). The cells were clamped using the patch clamp system and a whole-cell voltage-clamp mode was executed to elicit hERG currents at corresponding voltages. Both cisapride (a positive control from sigma) and test compounds were applied to the cells in serially diluted concentrations. Tail currents of the hERG channels were meticulously recorded using Patchcontrol HT and Patchmaster to obtain the peak tail current at each concentration. The current recorded in a non-compound containing solution served as a control for each cell. For each cell, the current was recorded independently twice, and at least two cells were recorded for each concentration.

**Synthesis**

The target compound **1-9** were synthesized according to the general procedure outlined in Scheme 1. At first, the benzoic acid intermediate was prepared. The synthetic route initiates with a palladium-catalyzed Buchwald-Hartwig amination reaction form the corresponding aniline derivative. The aniline is then subjected to a base-mediated hydrolysis, yielding the benzoic acid intermediate. Secondly, the intermediate of aryl cyclopentylamines was synthesized through a Grignard reaction involving ethyl magnesium bromide. This reaction transforms an aryl nitrile into an intermediate featuring a primary amine group, setting the stage for subsequent chemical modifications. Finally, the carboxylic acid intermediate **1d** was condensed with the amine intermediate through HATU activation to generate the final products. To synthesize compound **4**, **8** and **9**, the process starts with a Boc-protected amine to maintain the selectivity during the amide condensation reaction. This step leads to a Boc-protected amide intermediate. The pathway concludes with the deprotection of the amine group using TFA, yielding the final amine product.

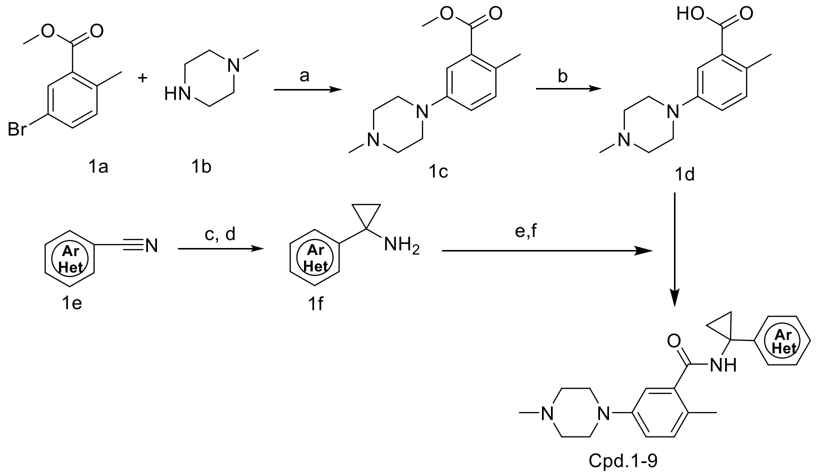

**Scheme1**. Reagents and conditions: (a) $Pd_2(dba)_3$, X-Phos, $Cs_2CO_3$, toluene, 110 °C; (b) NaOH, THF, EtOH, $H_2O$, 25 °C; (c) EtMgBr, $Ti(OEt)_4$, THF, -78 °C - 25 °C; (d) $BF_3 \cdot Et_2O$, 25 °C; (e) HATU, DIPEA, DMF, 50 °C; (f) TFA, DCM, 25 °C.

**Procedure for the synthesis of compound 1c**

Under the inert atmosphere created by nitrogen gas, methyl 5-bromo-2-methylbenzoate (2.98 g, 13 mmol, 1 eq.), 1-methylpiperazine (1.30 g, 13 mmol, 1 eq.), $Pd_2(dba)_3$ (238.1 mg, 0.26 mmol, 0.02 eq.), XPhos (247.9 mg, 0.52 mmol, 0.04 eq.), $Cs_2CO_3$ (8.47 g, 26 mmol, 2 eq.), and 1,4-dioxane (40 mL) were sequentially added to a round-bottom flask. The resulting mixture was stirred at 110 °C for 16 hours. Once the reaction was completed, the reaction mixture was cooled to room temperature and filtered. The filtrate was then concentrated at a reduced pressure and dried to obtain the crude product, which was further purified by column chromatography to afford compound **1c** (2.5 g) as yellow solid.

**Procedure and characterization data for the synthesis of compound 1d**

The intermediate **1c** (4.69 g, 20 mmol, 1 eq.), NaOH (4.1 g, 10 mmol, 5 eq.), and a mixture of THF/MeOH/$H_2O$ (48 mL:16 mL:16 mL) were sequentially added to a flask. The resulting mixture was stirred at 50 °C for 2 hours. After the completion of the reaction, the mixture was cooled to room temperature, then the pH was adjusted to 5-6 by slowly adding dilute hydrochloric acid at 0 °C. After 10 minutes of standing, a white solid, that is compound **1d** (3.3 g), was precipitated from the mixture.

**2-methyl-5-(4-methylpiperazin-1-yl)benzoic acid (1d)**. White solid, 70% yield. $^1$H NMR (600 MHz, DMSO-$d_6$) δ 7.36 (d, J = 2.8 Hz, 1H), 7.16 (d, J = 8.5 Hz, 1H), 7.08 (dd, J = 8.4, 2.8 Hz, 1H), 2.97 (s, 4H), 2.56 (s, 3H), 2.41 (s, 3H). LC-MS (ESI, m/z): $C_{13}H_{18}N_2O_2$, [M+H]+= 235.19.

**Procedure for the synthesis of compound 1f**

Under the inert atmosphere created by nitrogen gas, a solution of the reagent **1e** (1 equiv., 2 mmol,) in THF (15 mL) was cooled to -78 °C. Then, $Ti(OEt)_4$ (1.1 equiv, 2.2 mmol) was added to the solution, followed by the slow addition of ethylmagnesium bromide (2.2 equiv., 4.4 mmol, 3 M in Et2O, 1.47 mL). After this operation, the reaction mixture was stirred at room temperature for 1 hour. Then, $BF_3 \cdot Et_2O$ (2 equiv, 4 mmol) was added at room temperature and the mixture was stirred for an additional 1 hour. Once the reaction was completed, the reaction mixture was quenched at 0 °C by adding saturated ammonium chloride solution (5 mL). Extraction was then performed using dichloromethane and water. The organic layer was separated and dried over anhydrous $Na_2SO_4$, followed by concentration at a reduced pressure to remove the organic solvent. The crude product was purified by preparative liquid chromatography to obtain compound **1f**.

**General procedure and characterization data for the synthesis of compound 1-9**

HATU (1.2 equiv., 0.24 mmol) and DIPEA (3.6 equiv., 0.72 mmol) were added to a solution of amine (1 equiv., 0.2 mmol) and carboxylic acid (1.2 equiv., 0.24 mmol) in DMF. The reaction mixture was stirred at 50 °C for 3 hours. Upon completion, as indicated by TLC, the reaction mixture was diluted with water, extracted with ethyl acetate, and dried over anhydrous $Na_2SO_4$. The solvent was removed at a reduced pressure to yield the crude product. For the compounds with a BOC group, the crude product was treated with a mixture of DCM/TFA (v/v = 2:1) and stirred at room temperature for 3 hours. After the reaction was complete, as indicated by TLC, the mixture was diluted with ethyl acetate (25 mL), washed with water (20 mL), neutralized with $NaHCO_3$, dried over anhydrous $Na_2SO_4$, and evaporated in vacuo. The crude residue was then purified by preparative liquid chromatography. For compounds without a BOC group, the deprotection step with TFA is not necessary, and the product from the amide coupling reaction can be directly purified by preparative liquid chromatography.

**N-(1-(8-chloroquinolin-4-yl)cyclopropyl)-2-methyl-5-(4-methylpiperazin-1-yl)benzamide (1).** Light yellow solid, 68% yield. $^1$H NMR (600 MHz, DMSO-$d_6$) δ 9.19 (s, 1H), 8.99 (d, J = 4.3 Hz, 1H), 8.65 (d, J = 8.3 Hz, 1H), 7.97 (d, J = 7.1 Hz, 1H), 7.81 (d, J = 4.3 Hz, 1H), 7.64 (t, J = 8.0 Hz, 1H), 7.01 (d, J = 8.4 Hz, 1H), 6.90 (dd, J = 8.3, 2.4 Hz, 1H), 6.64 (d, J = 2.3 Hz, 1H), 3.06 (s, 4H), 2.71 – 2.60 (m, 3H), 2.50 (s, 3H), 1.99 (dq, J = 12.0, 6.9, 6.4 Hz, 1H), 1.90 (s, 3H), 1.39 (t, J = 5.4 Hz, 2H), 1.29 (d, J = 8.5 Hz, 2H). $^{13}$C NMR (150 MHz, DMSO-d6) δ 170.29, 151.38, 147.63, 144.59, 137.82, 133.67, 131.46, 129.61, 129.03, 126.82, 125.32, 124.13, 117.44, 114.86, 53.35, 46.96, 34.08, 29.50, 29.45, 18.33, 14.19. LC-MS (ESI, m/z): $C_{25}H_{27}ClN_4O$, [M+H]+= 435.1981.

**2-methyl-5-(4-methylpiperazin-1-yl)-N-(1-(quinolin-4-yl)cyclopropyl)benzamide formate (2).** White solid, 52% yield. $^1$H NMR (600 MHz, DMSO-$d_6$) δ 8.65 (s, 1H), 8.35 (d, J = 4.4 Hz, 1H), 8.15 (d, J = 8.4 Hz, 1H), 7.65 (s, 1H), 7.53 (d, J = 8.4 Hz, 1H), 7.24 (t, J = 7.6 Hz, 1H), 7.17 (d, J = 4.4 Hz, 1H), 7.13 (t, J = 7.6 Hz, 1H), 6.44 (d, J = 8.4 Hz, 1H), 6.32 (dd, J = 8.5, 2.7 Hz, 1H), 6.08 (d, J = 2.7 Hz, 1H), 1.98 (d, J = 3.2 Hz, 4H), 1.94 (t, J = 4.9 Hz, 4H), 1.72 (s, 3H), 1.37 (s, 3H), 0.85 (q, J = 5.0 Hz, 2H), 0.73 (q, J = 5.0 Hz, 2H). $^{13}$C NMR (151 MHz, DMSO) δ 170.39, 163.90, 150.65, 148.97, 148.74, 146.90, 137.66, 131.33, 130.13, 129.36, 127.43, 126.59, 125.88, 125.43, 122.94, 116.95, 114.49, 54.80, 48.49, 45.92, 34.02, 18.35, 14.00. LC-MS (ESI, m/z): $C_{25}H_{28}N_4O$, [M+H]+=401.2349.

**N-(1-(1H-benzo[d]imidazol-4-yl)cyclopropyl)-2-methyl-5-(4-methylpiperazin-1-yl)benzamide formate (3)**. White solid, 43% yield. $^1$H NMR (600 MHz, DMSO-$d_6$) δ 8.97 (s, 1H), 8.13 (s, 1H), 7.88 (d, J = 8.0 Hz, 1H), 7.45 (d, J = 7.8 Hz, 1H), 7.40 (t, J = 7.9 Hz, 1H), 7.27 (d, J = 8.6 Hz, 1H), 7.18 (s, 1H), 7.14 (dd, J = 8.7, 2.8 Hz, 1H), 7.05 (d, J = 8.5 Hz, 1H), 7.00 (s, 1H), 6.90 (dd, J = 8.2, 2.6 Hz, 1H), 3.15 – 3.11 (m, 4H), 2.45 – 2.42 (m, 4H), 2.22 (s, 2H), 2.20 (s, 3H), 2.18 (s, 1H), 1.30 (q, J = 4.8 Hz, 2H), 1.24 (d, J = 5.4 Hz, 2H). LC-MS (ESI, m/z): $C_{23}H_{27}N_5O$, [M+H]+=390.2299.

**N-(1-(1H-indol-3-yl)cyclopropyl)-2-methyl-5-(4-methylpiperazin-1-yl)benzamide formate (4).** White solid, 51% yield. $^1$H NMR (600 MHz, DMSO-$d_6$) δ 10.79 (s, 1H), 8.88 (s, 1H), 8.14 (s, 1H), 7.89 (d, J = 8.0 Hz, 1H), 7.32 (d, J = 8.1 Hz, 1H), 7.21 (d, J = 2.4 Hz, 1H), 7.06 (t, J = 7.5 Hz, 1H), 7.02 – 6.92 (m, 2H), 6.85 (dd, J = 8.4, 2.6 Hz, 1H), 6.73 (d, J = 2.6 Hz, 1H), 3.05 (t, J = 5.0 Hz, 4H), 2.44 (t, J = 4.9 Hz, 4H), 2.22 (s, 3H), 2.07 (s, 3H), 1.23 (s, 2H), 1.13 (s, 2H). LC-MS (ESI, m/z): $C_{24}H_{28}N_4O$, [M+H]+=389.2342.

**N-(1-(benzofuran-2-yl)cyclopropyl)-2-methyl-5-(4-methylpiperazin-1-yl)benzamide formate (5)**. White solid, 39% yield. $^1$H NMR (600 MHz, DMSO-$d_6$) δ 9.04 (s, 1H), 8.13 (s, 1H), 7.56 – 7.51 (m, 1H), 7.48 – 7.44 (m, 1H), 7.23 – 7.18 (m, 2H), 7.10 (d, J = 8.3 Hz, 1H), 6.98 (d, J = 2.6 Hz, 1H), 6.95 (dd, J = 8.3, 2.7 Hz, 1H), 6.66 (s, 1H), 3.23 (s, 4H), 2.83 (s, 4H), 2.25 (s, 3H), 2.00 (dt, J = 12.3, 6.9 Hz, 1H), 1.47 (q, J = 5.0 Hz, 2H), 1.30 (q, J = 5.0 Hz, 2H), 1.24 (d, J = 5.0 Hz, 2H). LC-MS (ESI, m/z): $C_{24}H_{27}N_3O_2$, [M+H]+=390.2188.

**N-(1-(benzo[b]thiophen-3-yl)cyclopropyl)-2-methyl-5-(4-methylpiperazin-1-yl)benzamide formate (6)**. White solid, 44% yield. $^1$H NMR (600 MHz, DMSO-$d_6$) δ 9.04 (s, 1H), 8.33 (d, J = 7.9 Hz, 1H), 8.14 (s, 1H), 7.95 (d, J = 7.9 Hz, 1H), 7.65 (s, 1H), 7.39 (dt, J = 23.1, 7.3 Hz, 2H),

6.98 (d, J = 8.3 Hz, 1H), 6.85 (dd, J = 8.5, 2.7 Hz, 1H), 6.66 (d, J = 2.7 Hz, 1H), 3.04 (t, J = 5.0 Hz, 4H), 2.45 (s, 3H), 2.23 (s, 2H), 1.99 (s, 3H), 1.26 – 1.20 (m, 4H), 1.17 (d, J = 5.1 Hz, 2H). LC-MS (ESI, m/z): $C_{24}H_{27}N_3OS$, [M+H]+=406.2122.

**2-methyl-N-(1-(1-methyl-1H-indol-4-yl)cyclopropyl)-5-(4-methylpiperazin-1-yl)benzamide (7)**. White solid, 22% yield. $^1$H NMR (600 MHz, DMSO-$d_6$) δ 8.96 (s, 1H), 7.32 – 7.29 (m, 2H), 7.12 (d, J = 7.1 Hz, 1H), 7.06 (t, J = 7.6 Hz, 1H), 6.99 (d, J = 8.4 Hz, 1H), 6.88 – 6.84 (m, 2H), 6.71 (d, J = 2.4 Hz, 1H), 3.78 (s, 3H), 3.06 (s, 4H), 2.48 (s, 3H), 2.25 (s, 3H), 2.05 (s, 3H), 2.00 (dt, J = 20.0, 7.1 Hz, 1H), 1.19 (d, J = 5.0 Hz, 4H). $^{13}$C NMR (150 MHz, DMSO-d6) δ 169.89, 148.98, 138.15, 137.10, 134.61, 131.28, 130.13, 129.21, 127.52, 125.64, 120.87, 118.82, 116.83, 114.73, 108.92, 100.45, 54.90, 48.65, 34.91, 33.02, 31.76, 30.85, 29.49, 29.44, 22.57, 18.62, 14.65. LC-MS (ESI, m/z): $C_{25}H_{30}N_4O$, [M+H]+=403.2473.

**N-(1-(1H-pyrrolo[3,2-c]pyridin-4-yl)cyclopropyl)-2-methyl-5-(4-methylpiperazin-1-yl)benzamide formate (8)**. White solid, 40% yield. $^1$H NMR (600 MHz, DMSO-$d_6$) δ 11.43 (s, 1H), 9.21 (s, 1H), 8.18 (s, 1H), 8.00 (d, J = 5.6 Hz, 1H), 7.36 (s, 1H), 7.20 (d, J = 5.9 Hz, 1H), 7.06 (d, J = 8.1 Hz, 1H), 6.93 (d, J = 7.6 Hz, 1H), 6.82 (s, 1H), 6.66 (s, 1H), 3.14 (s, 3H), 2.54 (s, 3H), 2.25 (d, J = 19.6 Hz, 2H), 2.14 (s, 1H), 1.99 (dq, J = 13.0, 7.1, 6.7 Hz, 2H), 1.63 (s, 1H), 1.49 – 1.43 (m, 1H), 1.29 (d, J = 6.9 Hz, 2H), 1.21 (s, 2H), 0.85 (t, J = 6.9 Hz, 1H). LC-MS (ESI, m/z): $C_{24}H_{27}N_5O$, [M+H]+=390.2286.

**N-(1-(1H-indol-4-yl)cyclopropyl)-2-methyl-5-(4-methylpiperazin-1-yl)benzamide (9)**. White solid, 45% yield. $^1$H NMR (600 MHz, DMSO-$d_6$) δ 11.05 (s, 1H), 8.96 (s, 1H), 7.32 (s, 1H), 7.26 (d, J = 8.0 Hz, 1H), 7.07 (d, J = 7.3 Hz, 1H), 6.99 (d, J = 8.5 Hz, 2H), 6.88 (s, 1H), 6.85 (d, J = 8.5 Hz, 1H), 6.70 (s, 1H), 3.05 (d, J = 5.1 Hz, 3H), 2.43 (s, 3H), 2.21 (s, 2H), 2.05 (s, 2H), 1.98 (dt, J = 13.7, 7.6 Hz, 2H), 1.23 (s, 2H), 1.19 (s, 2H), 0.94 (t, J = 7.4 Hz, 1H), 0.85 (t, J = 6.8 Hz, 1H). LC-MS (ESI, m/z): $C_{24}H_{28}N_4O$, [M+H]+=389.2344.

The synthesis of **compound 10-36** was shown in Scheme 2. The synthesis of the aminopyridine intermediate starts with palladium-catalyzed coupling of bromopyridine with cyclopropyl cyanide, yielding compound **2b**. This is followed by cyano hydrolysis to form amide **2c**, which is subsequently transformed into aminopyridine **2d** through a Hofmann rearrangement. The aminopyridine intermediate was then condensed with the benzoic acid **2e** by adding HATU to facilitated the reaction to yield bromo-substituted compound **2f**. Next, **2g** underwent a palladium-catalyzed coupling with a Boc-protected amine, ensuring reaction selectivity and yielding the Boc-protected intermediate. The final step involved Boc deprotection to generate the final products. In a parallel route, direct palladium-catalyzed coupling of **2g** was conducted to generate the final products without necessitating a Boc deprotection step. Compound **19** was separately synthesized via a palladium-catalyzed Suzuki-Miyaura coupling reaction between the brominated intermediate and the boronic ester intermediate, yielding the desired product.

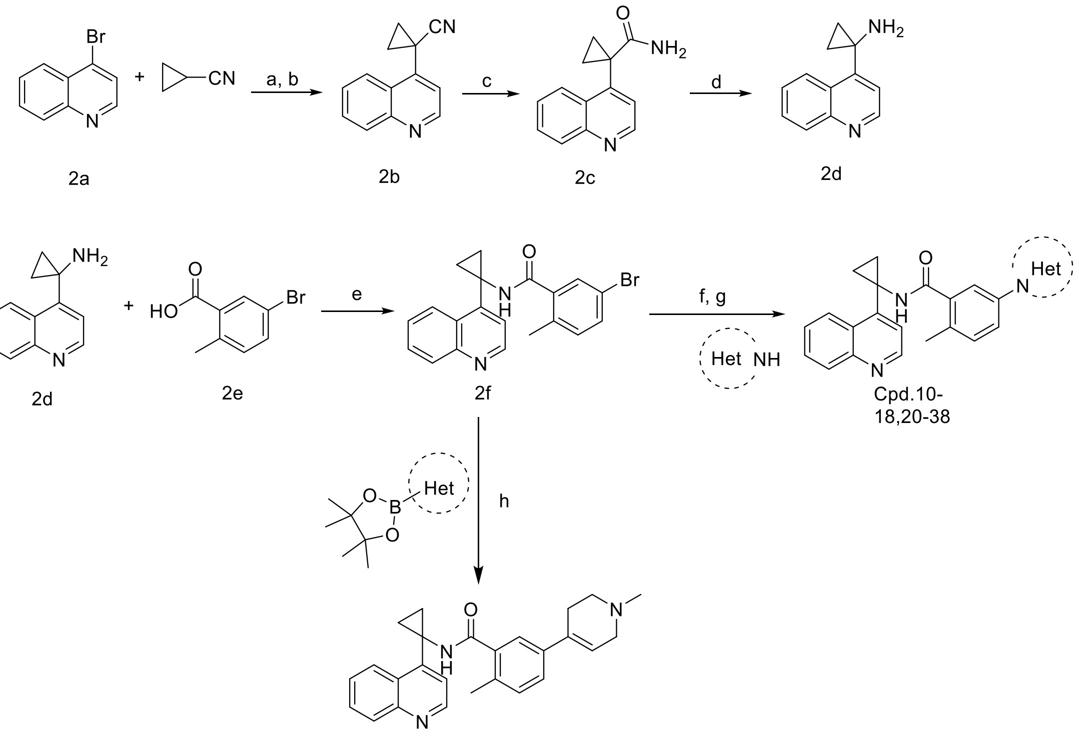

Cpd.19

**Scheme 2**. Reagents and conditions: Reagents and conditions: (a) $Pd_2dba_3$, NiXantphos, THF, 25 °C; (b) CPME, LiHMDS, 60 °C; (c) NaOH, *i*-PrOH, 90 °C; (d) NaClO, NaOH, *t*-BuOH, 0 °C - 25 °C. (e) HATU, DIPEA, DMF, 50 °C; (f) $Pd(OAc)_2$, Ru-Phos, $Cs_2CO_3$, 1,4-dioxane, 110 °C; (g) TFA, DCM, 25 °C. (h) $Pd_2(dba)_3$, XPhos, $Cs_2CO_3$, 1,4-dioxane, 100 °C.

**Procedure and characterization data for the synthesis of compound 2b**

Under the inert atmosphere created by nitrogen gas, $Pd_2(dba)_3$ (0.05 eq., 0.625 mmol, 582 mg) and NiXantphos (0.1 equiv., 1.25 mmol, 696 mg) were dissolved in THF (36 mL) and stirred at room temperature for 5 minutes. Subsequently, a solution of cyclopentyl methyl ether (36 mL), 4-bromoquinoline (1 equiv., 12.5 mmol, 2.6 g), and cyclopropanecarbonitrile (1.5 equiv., 18.75 mmol, 1.29 g) were added to the reaction mixture. LiHMDS (2 equiv., 25.8 mmol, 1 M in THF, 25.8 mL) was added dropwise. After this, the reaction mixture was stirred at 60 °C for 1 hour. At the end of the reaction, extraction was conducted using dichloromethane and water. The organic layer was separated. It was dried over anhydrous $Na_2SO_4$ and concentrated under a reduced pressure to remove the solvent. The crude product was purified by column chromatography, yielding 1.8 g yellow solid (MS m/z (ESI): 194.98 [M+H]+).

**Procedure and characterization data for the synthesis of compound 2c**

NaOH (5 equiv, 5 mmol, 200 mg) was added to a solution of compound **2b** (1 equiv., 1 mmol, 194.3 mg) in isopropanol (4 mL). The mixture was then stirred for 6 h until compound 2b consumed completely as indicated by TLC. Upon this, 181 mg brown solid was obtained by extracting the mixture with dichloromethane (DCM) and water and concentrating the mixture under a reduced pressure.

**1-(quinolin-4-yl)cyclopropane-1-carboxamide (2c).** Brown solid., 85% yield. $^1H$ NMR (600 MHz, DMSO-$d_6$) δ 8.85 (d, J = 4.4 Hz, 1H), 8.10 (dd, J = 8.4, 1.4 Hz, 1H), 8.05 (d, J = 8.4 Hz,

1H), 7.76 (ddd, J = 8.4, 6.8, 1.4 Hz, 1H), 7.65 (ddd, J = 8.3, 6.7, 1.3 Hz, 1H), 7.50 (d, J = 4.3 Hz, 1H), 7.02 (s, 1H), 6.48 (s, 1H), 1.60 (q, J = 3.3 Hz, 2H), 1.10 (q, J = 3.4 Hz, 2H). LC-MS (ESI, m/z): $C_{13}H_{12}N_2O$, [M+H]+= 213.15.

**Procedure and characterization data for the synthesis of compound 2d**

Under an atmosphere of nitrogen gas, a solution of **2c** (1 equiv., 0.5 mmol, 106.2 mg) in tert-butanol (15 mL) was cooled to 0 °C. 3 M NaOH (1.3 mL) and NaClO (1.6 mL) were added to the solution, followed by stirring the mixture at room temperature for 6 hours. After the reaction, the mixture was concentrated under a reduced pressure to obtain the crude product, which was further purified by column chromatography, yielding 45 mg compound **2d**.

**1-(quinolin-4-yl)cyclopropan-1-amine (2d).** Brown solid., 49% yield. $^1$H NMR (600 MHz, DMSO-$d_6$) δ 8.81 (d, J = 4.3 Hz, 1H), 8.50 (dd, J = 8.5, 1.4 Hz, 1H), 8.04 (dd, J = 8.4, 1.3 Hz, 1H), 7.76 (ddd, J = 8.4, 6.7, 1.4 Hz, 1H), 7.66 (ddd, J = 8.2, 6.7, 1.3 Hz, 1H), 7.42 (d, J = 4.3 Hz, 1H), 2.47 (s, 2H), 1.09 – 1.03 (m, 2H), 0.92 – 0.88 (m, 2H). LC-MS (ESI, m/z): $C_{12}H_{12}N_2$, [M+H]+= 185.18.

**Procedure and characterization data for the synthesis of compound 2f**

Compound **2d** (1equiv., 2.00 g, 10.86 mmol), **2e** (1.2 equiv., 2.80 g, 13.03 mmol), DIPEA (3 equiv., 4.21 g, 32.57 mmol), HATU (1.2 equiv., 4.96 g, 13.03 mmol) and DMF (15 mL) were added to the flask sequentially and reacted at 50 °C for 3 hours. After the reaction, the mixture was concentrated under a reduced pressure to obtain the crude product, which was further purified by column chromatography to obtain 2.1 g compound **2f** (MS m/z (ESI): 380.71 [M+H]+)

**General procedure and characterization data for the synthesis of compounds 10-18, 20-36**

The intermediate **2f** (1 equiv, 0.1 mmol) was added to a sealed tube, followed by adding tert-butyl 3,6-diaza-bicyclo[3.1.1]heptane-6-carboxylate (2 equiv, 0.2 mmol), $Cs_2CO_3$ (3 equiv, 0.3 mmol), $Pd(OAc)_2$ (0.1 equiv., 0.01 mmol), RuPhos (0.2 equiv., 0.02 mmol), and 1,4-dioxane (1.5 mL) sequentially. The reaction mixture was stirred under the atmosphere of nitrogen gas at 110 °C for 16 hours. The reaction mixture was then concentrated under a reduced pressure to afford crude product. For compounds with a BOC group, the crude product was treated with a mixture of DCM/4 M HCl (v/v = 1:1) and stirred at room temperature for 2 hours. Then, the crude product was purified by preparative liquid chromatography. Desired solid can be obtained by lyophilization of the purified product. For compounds without a BOC group, the deprotection step with TFA is not necessary, and the product from the amide coupling reaction can be directly purified by preparative liquid chromatography.

**2-methyl-5-(piperazin-1-yl)-N-(1-(quinolin-4-yl)cyclopropyl)benzamide formate (10)**. White solid, 55% yield. $^1$H NMR (600 MHz, DMSO-$d_6$) δ 9.19 (s, 1H), 8.87 (d, J = 4.4 Hz, 1H), 8.67 (dd, J = 8.5, 1.4 Hz, 1H), 8.26 (s, 1H), 8.05 (dd, J = 8.5, 1.3 Hz, 1H), 7.76 (ddd, J = 8.3, 6.7, 1.4 Hz, 1H), 7.69 (d, J = 4.4 Hz, 1H), 7.65 (ddd, J = 8.2, 6.8, 1.3 Hz, 1H), 6.98 (d, J = 8.4 Hz, 1H), 6.86 (dd, J = 8.4, 2.7 Hz, 1H), 6.62 (d, J = 2.6 Hz, 1H), 3.11 – 3.05 (m, 4H), 2.98 (t, J = 4.9 Hz, 4H), 1.90 (s, 3H), 1.38 (q, J = 5.0, 4.6 Hz, 2H), 1.27 – 1.22 (m, 2H). LC-MS (ESI, m/z): $C_{24}H_{26}N_4O$, [M+H]+= 387.2168.

**5-(4-(2-hydroxy-2-methylpropyl)piperazin-1-yl)-2-methyl-N-(1-(quinolin-4-yl)cyclopropyl)benzamide formate (11).** White solid, 15% yield. $^{1}H$ NMR (600 MHz, DMSO-$d_6$) δ 9.15 (s, 1H), 8.87 (d, J = 4.4 Hz, 1H), 8.67 (dd, J = 8.4, 1.4 Hz, 1H), 8.17 (s, 1H), 8.05 (dd, J = 8.5, 1.3 Hz, 1H), 7.76 (ddd, J = 8.3, 6.8, 1.4 Hz, 1H), 7.69 (d, J = 4.4 Hz, 1H), 7.64 (ddd, J = 8.3, 6.8, 1.3 Hz, 1H), 6.95 (d, J = 8.4 Hz, 1H), 6.82 (dd, J = 8.4, 2.7 Hz, 1H), 6.59 (d, J = 2.7 Hz, 1H), 2.99 (t, J = 4.9 Hz, 4H), 2.61 (t, J = 4.9 Hz, 4H), 2.22 (s, 2H), 1.89 (s, 3H), 1.41 – 1.33 (m, 2H), 1.25 – 1.23 (m, 2H), 1.10 (s, 6H). LC-MS (ESI, m/z): $C_{28}H_{34}N_4O_2$, [M+H]+=459.2761.

**5-(4-(2-(dimethylamino)ethyl)piperazin-1-yl)-2-methyl-N-(1-(quinolin-4-yl)cyclopropyl)benzamide formate (12).** White solid, 31% yield. $^{1}H$ NMR (600 MHz, DMSO-$d_6$) δ 9.91 (s, 1H), 8.86 (d, J = 4.4 Hz, 1H), 8.28 – 8.25 (m, 1H), 8.24 (s, 1H), 8.03 (d, J = 8.4 Hz, 1H), 7.75 (ddd, J = 8.3, 6.7, 1.4 Hz, 1H), 7.59 (ddd, J = 8.3, 6.7, 1.3 Hz, 1H), 7.45 (d, J = 4.4 Hz, 1H), 7.06 (d, J = 8.2 Hz, 1H), 6.96 – 6.89 (m, 2H), 5.75 (q, J = 6.8 Hz, 1H), 3.10 (t, J = 4.8 Hz, 4H), 2.56 (q, J = 5.8, 4.5 Hz, 6H), 2.49 – 2.46 (m, 2H), 2.29 (s, 6H), 2.17 (s, 3H), 1.89 (d, J = 6.9 Hz, 3H). LC-MS (ESI, m/z): $C_{28}H_{35}N_5O$, [M+H]+=458.2886.

**2-methyl-5-(4-(methylsulfonyl)piperazin-1-yl)-N-(1-(quinolin-4-yl)cyclopropyl)benzamide (13)**. White solid, 30% yield. $^{1}H$ NMR (600 MHz, DMSO-$d_6$) δ 9.18 (s, 1H), 8.87 (d, J = 4.4 Hz, 1H), 8.66 (dd, J = 8.5, 1.4 Hz, 1H), 8.05 (dd, J = 8.5, 1.3 Hz, 1H), 7.76 (ddd, J = 8.3, 6.7, 1.4 Hz, 1H), 7.70 (d, J = 4.4 Hz, 1H), 7.66 (ddd, J = 8.3, 6.8, 1.3 Hz, 1H), 7.00 (d, J = 8.3 Hz, 1H), 6.89 (dd, J = 8.4, 2.7 Hz, 1H), 6.65 (d, J = 2.6 Hz, 1H), 3.20 (dd, J = 6.4, 3.5 Hz, 4H), 3.13 (dd, J = 6.5, 3.5 Hz, 4H), 2.91 (s, 3H), 1.90 (s, 3H), 1.38 (q, J = 5.0, 4.6 Hz, 2H), 1.27 – 1.23 (m, 2H). LC-MS (ESI, m/z): $C_{25}H_{28}N_4O_3S$, [M+H]+=465.1963.

**2-methyl-5-(3-methylpiperazin-1-yl)-N-(1-(quinolin-4-yl)cyclopropyl)benzamide formate (14)**. White solid, 42% yield. $^{1}H$ NMR (600 MHz, DMSO-$d_6$) δ 9.18 (s, 1H), 8.87 (d, J = 4.3 Hz, 1H), 8.67 (d, J = 8.4 Hz, 1H), 8.28 (s, 1H), 8.05 (d, J = 8.4 Hz, 1H), 7.76 (t, J = 7.6 Hz, 1H), 7.69 (d, J = 4.4 Hz, 1H), 7.65 (t, J = 7.6 Hz, 1H), 6.97 (d, J = 8.4 Hz, 1H), 6.85 (dd, J = 8.4, 2.4 Hz, 1H), 6.60 (d, J = 2.3 Hz, 1H), 3.48 – 3.39 (m, 2H), 3.05 (d, J = 11.7 Hz, 1H), 2.91 (s, 1H), 2.83 (t, J = 11.1 Hz, 1H), 2.56 (t, J = 11.4 Hz, 1H), 2.25 (t, J = 11.0 Hz, 1H), 1.90 (s, 3H), 1.37 (t, J = 5.4 Hz, 2H), 1.25 (t, J = 5.4 Hz, 2H), 1.08 (d, J = 6.3 Hz, 3H). LC-MS (ESI, m/z): $C_{25}H_{28}N_4O$, [M+H]+=401.2338.

**2-methyl-5-(2-methylpiperazin-1-yl)-N-(1-(quinolin-4-yl)cyclopropyl)benzamide formate (15)**. White solid, 41% yield. $^{1}H$ NMR (600 MHz, DMSO-$d_6$) δ 9.90 (s, 1H), 8.86 (d, J = 4.4 Hz, 1H), 8.31 – 8.20 (m, 2H), 8.03 (d, J = 8.4 Hz, 1H), 7.75 (ddd, J = 8.3, 6.8, 1.4 Hz, 1H), 7.59 (ddd, J = 8.4, 6.8, 1.3 Hz, 1H), 7.44 (d, J = 4.4 Hz, 1H), 7.10 – 7.02 (m, 1H), 6.90 (dt, J = 5.1, 2.2 Hz, 2H), 5.76 (q, J = 6.8 Hz, 1H), 3.83 (dt, J = 7.0, 3.6 Hz, 1H), 3.16 (dt, J = 12.1, 3.5 Hz, 1H), 3.08 – 3.02 (m, 1H), 2.99 (dd, J = 12.2, 3.6 Hz, 1H), 2.91 (td, J = 11.9, 11.3, 3.1 Hz, 1H), 2.87 – 2.76 (m, 2H), 2.18 (s, 3H), 1.89 (d, J = 6.9 Hz, 3H), 0.97 (d, J = 6.5 Hz, 3H). LC-MS (ESI, m/z): $C_{25}H_{28}N_4O$, [M+H]+=401.2332.

**5-(3,6-diazabicyclo[3.1.1]heptan-3-yl)-2-methyl-N-(1-(quinolin-4-yl)cyclopropyl)benzamide formate (16)**. White solid, 24% yield. $^{1}H$ NMR (600 MHz, DMSO-$d_6$) δ 9.17 (s, 1H), 8.87 (d, J = 4.4 Hz, 1H), 8.68 (dd, J = 8.4, 1.4 Hz, 1H), 8.30 (s, 1H), 8.05 (dd, J = 8.4, 1.2 Hz, 1H), 7.76 (ddd, J = 8.3, 6.8, 1.4 Hz, 1H), 7.69 (d, J = 4.4 Hz, 1H), 7.66 (ddd, J = 8.3, 6.8, 1.3 Hz, 1H), 6.98 (d, J =

8.4 Hz, 1H), 6.62 (dd, J = 8.4, 2.7 Hz, 1H), 6.40 (d, J = 2.7 Hz, 1H), 3.97 (d, J = 6.0 Hz, 2H), 3.49 (d, J = 11.0 Hz, 2H), 3.38 (d, J = 11.0 Hz, 2H), 2.64 (d, J = 7.6 Hz, 1H), 2.18 (s, 1H), 1.92 (s, 3H), 1.56 (d, J = 8.9 Hz, 1H), 1.37 (q, J = 5.0 Hz, 2H), 1.28 – 1.22 (m, 2H). $^{13}$C NMR (151 MHz, DMSO) δ 170.67, 165.06, 150.64, 148.75, 146.94, 146.66, 137.80, 131.42, 130.13, 129.36, 127.46, 126.61, 125.89, 122.92, 122.24, 111.52, 109.29, 56.29, 50.01, 34.05, 30.42, 18.29, 14.02. LC-MS (ESI, m/z): $C_{25}H_{26}N_4O$, [M+H]+=399.2191.

**5-(3,8-diazabicyclo[3.2.1]octan-8-yl)-2-methyl-N-(1-(quinolin-4-yl)cyclopropyl)benzamide formate (17)**. White solid, 51% yield. $^1$H NMR (600 MHz, DMSO-$d_6$) δ 9.16 (d, J = 2.5 Hz, 1H), 8.87 (d, J = 4.3 Hz, 1H), 8.66 (dd, J = 8.5, 1.4 Hz, 1H), 8.33 – 8.22 (m, 1H), 8.04 (dd, J = 8.4, 1.2 Hz, 1H), 7.76 (ddd, J = 8.3, 6.7, 1.4 Hz, 1H), 7.69 (dd, J = 4.5, 2.0 Hz, 1H), 7.64 (ddd, J = 8.3, 6.8, 1.4 Hz, 1H), 6.96 – 6.90 (m, 1H), 6.77 – 6.68 (m, 1H), 6.47 (q, J = 3.0 Hz, 1H), 4.09 – 3.99 (m, 2H), 2.88 (d, J = 12.0 Hz, 2H), 2.56 – 2.51 (m, 2H), 1.88 (s, 7H), 1.37 (q, J = 4.9, 4.5 Hz, 2H), 1.28 – 1.21 (m, 2H). LC-MS (ESI, m/z): $C_{26}H_{28}N_4O$, [M+H]+=413.2342.

**5-(hexahydropyrrolo[1,2-a]pyrazin-2(1H)-yl)-2-methyl-N-(1-(quinolin-4-yl)cyclopropyl)benzamide formate (18).** White solid, 41% yield. $^1$H NMR (600 MHz, DMSO-$d_6$) δ 9.16 (s, 1H), 8.87 (d, J = 4.4 Hz, 1H), 8.68 (dd, J = 8.5, 1.4 Hz, 1H), 8.16 (s, 1H), 8.05 (dd, J = 8.5, 1.2 Hz, 1H), 7.76 (ddd, J = 8.3, 6.7, 1.4 Hz, 1H), 7.68 (d, J = 4.4 Hz, 1H), 7.64 (ddd, J = 8.3, 6.8, 1.3 Hz, 1H), 6.95 (d, J = 8.4 Hz, 1H), 6.85 (dd, J = 8.4, 2.7 Hz, 1H), 6.59 (d, J = 2.7 Hz, 1H), 3.61 (dt, J = 10.8, 2.5 Hz, 1H), 3.52 – 3.47 (m, 1H), 3.05 – 2.97 (m, 2H), 2.61 (td, J = 11.6, 3.3 Hz, 1H), 2.27 (t, J = 10.6 Hz, 1H), 2.17 (td, J = 11.2, 3.2 Hz, 1H), 2.09 – 1.96 (m, 2H), 1.90 (s, 3H), 1.81 (ddt, J = 9.7, 6.3, 4.8 Hz, 1H), 1.75 – 1.64 (m, 2H), 1.40 – 1.30 (m, 3H), 1.26 – 1.23 (m, 2H). LC-MS (ESI, m/z): $C_{27}H_{30}N_4O$, [M+H]+=427.2496.

**5-(4-aminopiperidin-1-yl)-2-methyl-N-(1-(quinolin-4-yl)cyclopropyl)benzamide formate (20).** White solid, 44% yield. $^1$H NMR (600 MHz, DMSO-$d_6$) δ 9.20 (s, 1H), 8.87 (d, J = 4.4 Hz, 1H), 8.68 (d, J = 8.4 Hz, 1H), 8.42 (s, 1H), 8.05 (d, J = 8.3 Hz, 1H), 7.76 (ddd, J = 8.3, 6.7, 1.4 Hz, 1H), 7.69 (d, J = 4.4 Hz, 1H), 7.65 (ddd, J = 8.2, 6.7, 1.3 Hz, 1H), 6.95 (d, J = 8.4 Hz, 1H), 6.84 (dd, J = 8.4, 2.7 Hz, 1H), 6.61 (d, J = 2.6 Hz, 1H), 3.55 (d, J = 12.6 Hz, 2H), 2.98 (d, J = 11.6 Hz, 1H), 2.63 (td, J = 12.4, 2.4 Hz, 2H), 1.90 (s, 3H), 1.86 (d, J = 12.3 Hz, 2H), 1.47 (qd, J = 12.0, 3.9 Hz, 2H), 1.37 (q, J = 5.0 Hz, 2H), 1.28 – 1.20 (m, 2H). LC-MS (ESI, m/z): $C_{25}H_{28}N_4O$, [M+H]+=401.2346.

**5-(3-(dimethylamino)pyrrolidin-1-yl)-2-methyl-N-(1-(quinolin-4-yl)cyclopropyl)benzamide formate (21).** White solid, 33% yield. $^1$H NMR (600 MHz, DMSO-$d_6$) δ 9.13 (s, 1H), 8.87 (d, J = 4.4 Hz, 1H), 8.68 (dd, J = 8.5, 1.4 Hz, 1H), 8.19 (s, 1H), 8.05 (dd, J = 8.5, 1.2 Hz, 1H), 7.75 (ddd, J = 8.3, 6.8, 1.4 Hz, 1H), 7.68 (d, J = 4.4 Hz, 1H), 7.64 (ddd, J = 8.3, 6.8, 1.3 Hz, 1H), 6.90 (d, J = 8.3 Hz, 1H), 6.44 (dd, J = 8.3, 2.7 Hz, 1H), 6.21 (d, J = 2.6 Hz, 1H), 3.30 (dd, J = 9.1, 7.2 Hz, 1H), 3.24 (td, J = 9.1, 2.3 Hz, 1H), 3.12 (td, J = 9.4, 6.8 Hz, 1H), 2.91 (dd, J = 9.1, 7.8 Hz, 1H), 2.74 (dq, J = 9.6, 7.2 Hz, 1H), 2.18 (s, 6H), 2.11 (dtd, J = 11.9, 6.7, 2.3 Hz, 1H), 1.89 (s, 3H), 1.75 (dq, J = 11.9, 9.2 Hz, 1H), 1.40 – 1.34 (m, 2H), 1.27 – 1.22 (m, 2H). LC-MS (ESI, m/z): C26H30N4O, [M+H]+=415.2504.

**2-methyl-5-(3-(methylamino)pyrrolidin-1-yl)-N-(1-(quinolin-4-yl)cyclopropyl)benzamide formate (22)**. White solid, 18% yield. $^1$H NMR (600 MHz, DMSO-$d_6$) δ 9.15 (s, 1H), 8.87 (d, J =

4.4 Hz, 1H), 8.68 (dd, J = 8.6, 1.4 Hz, 1H), 8.32 (s, 1H), 8.05 (d, J = 8.4 Hz, 1H), 7.76 (ddd, J = 8.4, 6.7, 1.4 Hz, 1H), 7.69 (d, J = 4.4 Hz, 1H), 7.65 (ddd, J = 8.2, 6.6, 1.3 Hz, 1H), 6.91 (d, J = 8.3 Hz, 1H), 6.44 (dd, J = 8.3, 2.6 Hz, 1H), 6.23 (d, J = 2.6 Hz, 1H), 3.41 (p, J = 5.8 Hz, 1H), 3.34 (dd, J = 9.7, 6.5 Hz, 1H), 3.24 (td, J = 8.4, 5.7 Hz, 1H), 3.12 (td, J = 8.4, 6.3 Hz, 1H), 3.01 (dd, J = 9.8, 4.8 Hz, 1H), 2.38 (s, 3H), 2.13 (dq, J = 13.0, 6.5 Hz, 1H), 1.89 (s, 4H), 1.37 (q, J = 4.9 Hz, 2H), 1.26 – 1.22 (m, 2H). LC-MS (ESI, m/z): $C_{25}H_{28}N_4O$, [M+H]+=401.2349.

**2-methyl-N-(1-(quinolin-4-yl)cyclopropyl)-5-(2,7-diazaspiro[4.4]nonan-2-yl)benzamide formate (23)**. White solid, 25% yield. $^1$H NMR (600 MHz, DMSO-$d_6$) δ 9.13 (s, 1H), 8.87 (d, J = 4.4 Hz, 1H), 8.68 (dd, J = 8.5, 1.4 Hz, 1H), 8.24 (s, 1H), 8.05 (dd, J = 8.5, 1.3 Hz, 1H), 7.76 (ddd, J = 8.3, 6.8, 1.4 Hz, 1H), 7.70 – 7.62 (m, 2H), 6.90 (d, J = 8.3 Hz, 1H), 6.40 (dd, J = 8.3, 2.6 Hz, 1H), 6.18 (d, J = 2.6 Hz, 1H), 3.25 (d, J = 8.9 Hz, 1H), 3.19 (dd, J = 8.4, 5.2 Hz, 1H), 3.14 (d, J = 9.7 Hz, 1H), 3.09 (d, J = 9.6 Hz, 1H), 2.95 (ddt, J = 17.7, 11.0, 5.5 Hz, 2H), 2.02 – 1.98 (m, 1H), 1.93 (dt, J = 12.4, 7.4 Hz, 1H), 1.88 (s, 3H), 1.83 – 1.67 (m, 4H), 1.36 (d, J = 2.5 Hz, 1H), 1.25 – 1.23 (m, 3H). LC-MS (ESI, m/z): $C_{27}H_{30}N_4O$, [M+H]+=427.2521.

**5-(hexahydropyrrolo[3,4-c]pyrrol-2(1H)-yl)-2-methyl-N-(1-(quinolin-4-yl)cyclopropyl)benzamide formate (24)**. White solid, 35% yield. $^1$H NMR (600 MHz, DMSO-$d_6$) δ 9.14 (s, 1H), 8.87 (d, J = 4.3 Hz, 1H), 8.67 (d, J = 8.4 Hz, 1H), 8.31 (s, 1H), 8.05 (d, J = 8.4 Hz, 1H), 7.78 – 7.72 (m, 1H), 7.69 (d, J = 4.4 Hz, 1H), 7.65 (dd, J = 8.6, 6.6 Hz, 1H), 6.92 (d, J = 8.3 Hz, 1H), 6.54 (dd, J = 8.3, 2.6 Hz, 1H), 6.32 (d, J = 2.6 Hz, 1H), 3.20 (dd, J = 9.5, 7.1 Hz, 2H), 3.15 (dd, J = 11.4, 6.7 Hz, 2H), 3.03 (dd, J = 9.8, 2.9 Hz, 2H), 2.88 (td, J = 7.2, 3.7 Hz, 2H), 2.74 (dd, J = 11.4, 3.7 Hz, 2H), 1.88 (s, 3H), 1.36 (d, J = 2.2 Hz, 1H), 1.25 – 1.23 (m, 3H). LC-MS (ESI, m/z): $C_{26}H_{28}N_4O$, [M+H]+=413.2362.

**2-methyl-5-(1-methyloctahydro-6H-pyrrolo[3,4-b]pyridin-6-yl)-N-(1-(quinolin-4-yl)cyclopropyl)benzamide formate (25)**. White solid, 40% yield. $^1$H NMR (600 MHz, DMSO-$d_6$) δ 9.13 (s, 1H), 8.87 (d, J = 4.4 Hz, 1H), 8.69 (d, J = 8.4 Hz, 1H), 8.17 (s, 1H), 8.05 (d, J = 8.4 Hz, 1H), 7.75 (dd, J = 8.4, 6.8 Hz, 1H), 7.68 (d, J = 4.4 Hz, 1H), 7.64 (t, J = 7.6 Hz, 1H), 6.88 (d, J = 8.3 Hz, 1H), 6.38 (dd, J = 8.4, 2.6 Hz, 1H), 6.15 (d, J = 2.7 Hz, 1H), 3.29 (dd, J = 10.3, 2.7 Hz, 1H), 3.14 – 3.05 (m, 3H), 2.72 (q, J = 4.3 Hz, 1H), 2.62 (dt, J = 11.0, 4.3 Hz, 1H), 2.34 (td, J = 7.8, 4.0 Hz, 1H), 2.17 (s, 3H), 2.11 – 2.06 (m, 1H), 1.88 (s, 3H), 1.64 (dt, J = 8.5, 4.0 Hz, 1H), 1.56 (dq, J = 9.6, 4.8 Hz, 2H), 1.50 – 1.43 (m, 1H), 1.36 (q, J = 4.6 Hz, 2H), 1.25 (t, J = 3.5 Hz, 2H). LC-MS (ESI, m/z): $C_{28}H_{32}N_4O$, [M+H]+=441.2662.

**2-methyl-5-(6-methyl-2,6-diazaspiro[3.4]octan-2-yl)-N-(1-(quinolin-4-yl)cyclopropyl)benzamide formate (26)**. White solid, 54% yield. $^1$H NMR (600 MHz, DMSO-$d_6$) δ 9.13 (s, 1H), 8.86 (d, J = 4.4 Hz, 1H), 8.65 (dd, J = 8.5, 1.4 Hz, 1H), 8.17 (s, 1H), 8.04 (dd, J = 8.5, 1.3 Hz, 1H), 7.76 (ddd, J = 8.3, 6.8, 1.4 Hz, 1H), 7.68 (d, J = 4.4 Hz, 1H), 7.65 (ddd, J = 8.3, 6.8, 1.3 Hz, 1H), 6.91 (d, J = 8.2 Hz, 1H), 6.33 (dd, J = 8.2, 2.5 Hz, 1H), 6.12 (d, J = 2.5 Hz, 1H), 3.66 (d, J = 7.1 Hz, 2H), 3.59 (d, J = 7.1 Hz, 2H), 2.68 (s, 2H), 2.27 (s, 3H), 2.01 (t, J = 7.1 Hz, 2H), 1.87 (s, 3H), 1.42 – 1.32 (m, 2H), 1.26 – 1.22 (m, 4H). LC-MS (ESI, m/z): $C_{27}H_{30}N_4O$, [M+H]+=427.2497.

**5-(1,4-diazepan-1-yl)-2-methyl-N-(1-(quinolin-4-yl)cyclopropyl)benzamide formate (27).** White solid, 38% yield. $^1$H NMR (600 MHz, DMSO-$d_6$) δ 9.16 (s, 1H), 8.86 (d, J = 4.4 Hz, 1H),

8.68 (d, J = 8.4 Hz, 1H), 8.33 (s, 1H), 8.05 (d, J = 8.4 Hz, 1H), 7.76 (ddd, J = 8.3, 6.7, 1.4 Hz, 1H), 7.70 – 7.62 (m, 2H), 6.91 (d, J = 8.4 Hz, 1H), 6.63 (dd, J = 8.4, 2.8 Hz, 1H), 6.36 (d, J = 2.8 Hz, 1H), 3.47 (t, J = 4.9 Hz, 2H), 3.40 (t, J = 6.1 Hz, 2H), 2.93 (t, J = 4.8 Hz, 2H), 2.78 (t, J = 5.6 Hz, 2H), 1.89 (s, 3H), 1.83 (t, J = 5.8 Hz, 2H), 1.37 (q, J = 4.9, 4.5 Hz, 2H), 1.28 – 1.21 (m, 2H). LC-MS (ESI, m/z): $C_{25}H_{28}N_4O$, [M+H]+=401.2346.

**5-([1,3'-biazetidin]-1'-yl)-2-methyl-N-(1-(quinolin-4-yl)cyclopropyl)benzamide formate (28)**. White solid, 42% yield. $^1$H NMR (600 MHz, DMSO-$d_6$) δ 9.15 (s, 1H), 8.87 (d, J = 4.4 Hz, 1H), 8.66 (dd, J = 8.4, 1.3 Hz, 1H), 8.15 (s, 1H), 8.05 (d, J = 8.4 Hz, 1H), 7.76 (ddd, J = 8.3, 6.7, 1.4 Hz, 1H), 7.69 (d, J = 4.4 Hz, 1H), 7.65 (ddd, J = 8.3, 6.7, 1.3 Hz, 1H), 6.92 (d, J = 8.2 Hz, 1H), 6.34 (dd, J = 8.2, 2.6 Hz, 1H), 6.12 (d, J = 2.6 Hz, 1H), 3.74 (t, J = 7.3 Hz, 2H), 3.60 (t, J = 5.7 Hz, 1H), 3.51 (dd, J = 7.9, 4.5 Hz, 2H), 3.31 (t, J = 7.4 Hz, 4H), 2.03 (t, J = 7.2 Hz, 2H), 1.87 (s, 3H), 1.36 (t, J = 3.4 Hz, 2H), 1.26 – 1.23 (m, 2H). LC-MS (ESI, m/z): $C_{26}H_{28}N_4O$, [M+H]+=413.2365.

**5-([1,3'-bipyrrolidin]-1'-yl)-2-methyl-N-(1-(quinolin-4-yl)cyclopropyl)benzamide formate (29)**. White solid, 26% yield. $^1$H NMR (600 MHz, DMSO-$d_6$) δ 9.12 (s, 1H), 8.86 (d, J = 4.4 Hz, 1H), 8.68 (d, J = 8.4 Hz, 1H), 8.16 (s, 1H), 8.05 (d, J = 8.4 Hz, 1H), 7.75 (ddd, J = 8.4, 6.8, 1.4 Hz, 1H), 7.68 (d, J = 4.4 Hz, 1H), 7.64 (ddd, J = 8.2, 6.7, 1.3 Hz, 1H), 6.90 (d, J = 8.3 Hz, 1H), 6.43 (dd, J = 8.3, 2.6 Hz, 1H), 6.20 (d, J = 2.6 Hz, 1H), 3.29 (d, J = 9.1 Hz, 4H), 3.22 (td, J = 8.8, 3.0 Hz, 1H), 3.13 (td, J = 9.0, 6.9 Hz, 1H), 2.96 (dd, J = 9.1, 7.3 Hz, 1H), 2.79 (p, J = 7.0 Hz, 1H), 2.11 (dtd, J = 12.8, 6.7, 3.0 Hz, 1H), 1.99 (dq, J = 12.8, 7.0, 6.5 Hz, 1H), 1.88 (s, 3H), 1.83 (dq, J = 12.0, 8.7 Hz, 1H), 1.74 – 1.65 (m, 4H), 1.36 (t, J = 3.5 Hz, 2H), 1.25 – 1.24 (m, 2H). LC-MS (ESI, m/z): $C_{28}H_{32}N_4O$, [M+H]+=441.2659.

**5-(3-(4-hydroxy-4-methylpiperidin-1-yl)azetidin-1-yl)-2-methyl-N-(1-(quinolin-4-yl)cyclopropyl)benzamide formate (30)**. White solid, 21% yield. $^1$H NMR (600 MHz, DMSO-$d_6$) δ 9.14 (s, 1H), 8.87 (d, J = 4.3 Hz, 1H), 8.66 (d, J = 8.4 Hz, 1H), 8.14 (s, 1H), 8.05 (d, J = 8.4 Hz, 1H), 7.76 (t, J = 7.6 Hz, 1H), 7.69 (d, J = 4.4 Hz, 1H), 7.65 (t, J = 7.6 Hz, 1H), 6.93 (d, J = 8.2 Hz, 1H), 6.36 (d, J = 7.4 Hz, 1H), 6.15 (s, 1H), 3.86 (s, 2H), 3.48 (s, 2H), 3.31 (s, 5H), 1.88 (s, 3H), 1.47 (d, J = 18.4 Hz, 4H), 1.38 – 1.33 (m, 2H), 1.24 (t, J = 6.0 Hz, 3H), 1.11 (s, 3H). $^{13}$C NMR (151 MHz, DMSO) δ 169.28, 162.44, 149.57, 147.65, 145.78, 136.60, 130.09, 129.04, 128.28, 126.35, 125.53, 124.80, 121.90, 111.85, 109.41, 55.55, 53.80, 45.15, 39.81, 39.44, 32.88, 32.76, 17.30, 12.97. LC-MS (ESI, m/z): $C_{29}H_{34}N_4O_2$, [M+H]+=470.2760.

**5-(3-(3-hydroxypyrrolidin-1-yl)azetidin-1-yl)-2-methyl-N-(1-(quinolin-4-yl)cyclopropyl)benzamide formate (31)**. White solid, 28% yield. $^1$H NMR (600 MHz, DMSO-$d_6$) δ 9.15 (s, 1H), 8.87 (d, J = 4.4 Hz, 1H), 8.66 (dd, J = 8.5, 1.4 Hz, 1H), 8.14 (s, 1H), 8.05 (dd, J = 8.4, 1.3 Hz, 1H), 7.76 (ddd, J = 8.4, 6.8, 1.4 Hz, 1H), 7.69 (d, J = 4.4 Hz, 1H), 7.65 (ddd, J = 8.3, 6.8, 1.3 Hz, 1H), 6.91 (d, J = 8.2 Hz, 1H), 6.33 (dd, J = 8.2, 2.6 Hz, 1H), 6.12 (d, J = 2.5 Hz, 1H), 4.23 – 4.15 (m, 1H), 3.79 (t, J = 7.1 Hz, 2H), 3.52 (ddd, J = 10.1, 7.5, 5.2 Hz, 2H), 3.44 – 3.38 (m, 2H), 2.71 (dd, J = 9.6, 6.0 Hz, 1H), 2.57 (q, J = 7.8 Hz, 1H), 2.44 (td, J = 8.3, 5.3 Hz, 1H), 2.31 (dd, J = 9.7, 3.5 Hz, 1H), 1.96 (dq, J = 12.9, 7.4 Hz, 1H), 1.87 (s, 3H), 1.55 (dddd, J = 12.9, 8.0, 5.2, 3.3 Hz, 1H), 1.39 – 1.32 (m, 2H), 1.26 – 1.21 (m, 2H). $^{13}$C NMR (151 MHz, DMSO) δ 170.41, 163.59, 150.65, 149.94, 148.73, 146.89, 137.66, 131.15, 130.12, 129.36, 127.43, 126.59, 125.89, 123.10, 122.96, 112.74, 110.32, 69.83, 59.41, 56.58, 56.49, 53.43, 49.28, 34.76, 33.97, 18.36, 14.04. LC-MS (ESI, m/z): $C_{27}H_{30}N_4O_2$, [M+H]+=443.2455.

**5-(3-(4-fluoropiperidin-1-yl)azetidin-1-yl)-2-methyl-N-(1-(quinolin-4-yl)cyclopropyl)benzamide formate (32)**. White solid, 21% yield. $^{1}$H NMR (600 MHz, DMSO-$d_6$) δ 9.14 (s, 1H), 8.87 (d, J = 4.4 Hz, 1H), 8.66 (d, J = 8.4 Hz, 1H), 8.14 (s, 1H), 8.05 (d, J = 8.4 Hz, 1H), 7.76 (t, J = 7.6 Hz, 1H), 7.69 (d, J = 4.4 Hz, 1H), 7.65 (t, J = 7.6 Hz, 1H), 6.91 (d, J = 8.2 Hz, 1H), 6.34 (dd, J = 8.2, 2.5 Hz, 1H), 6.13 (d, J = 2.6 Hz, 1H), 4.69 (ddt, J = 49.0, 7.4, 3.6 Hz, 1H), 3.83 (t, J = 7.0 Hz, 2H), 3.43 (t, J = 6.5 Hz, 2H), 3.20 (t, J = 6.2 Hz, 1H), 2.40 (d, J = 10.2 Hz, 2H), 2.22 (d, J = 10.5 Hz, 2H), 1.87 (s, 5H), 1.74 – 1.64 (m, 2H), 1.36 (q, J = 5.0 Hz, 2H), 1.26 – 1.23 (m, 2H). LC-MS (ESI, m/z): $C_{28}H_{31}FN_4O$, [M+H]+=459.2573.

**5-(3-(3-hydroxypiperidin-1-yl)azetidin-1-yl)-2-methyl-N-(1-(quinolin-4-yl)cyclopropyl)benzamide formate (33)**. White solid, 23% yield. $^{1}$H NMR (600 MHz, DMSO-$d_6$) δ 9.14 (s, 1H), 8.87 (d, J = 4.4 Hz, 1H), 8.66 (dd, J = 8.5, 1.4 Hz, 1H), 8.15 (s, 1H), 8.05 (dd, J = 8.5, 1.3 Hz, 1H), 7.76 (ddd, J = 8.3, 6.7, 1.4 Hz, 1H), 7.69 (d, J = 4.4 Hz, 1H), 7.65 (ddd, J = 8.2, 6.7, 1.3 Hz, 1H), 6.91 (d, J = 8.2 Hz, 1H), 6.35 (dd, J = 8.2, 2.5 Hz, 1H), 6.14 (d, J = 2.5 Hz, 1H), 4.63 (s, 1H), 3.82 (q, J = 6.7 Hz, 2H), 3.42 (dt, J = 7.4, 5.7 Hz, 3H), 3.19 (p, J = 6.2 Hz, 1H), 2.71 (dd, J = 10.6, 4.1 Hz, 1H), 2.57 – 2.52 (m, 1H), 1.87 (s, 3H), 1.76 (tt, J = 11.8, 7.1 Hz, 2H), 1.67 – 1.59 (m, 2H), 1.41 – 1.32 (m, 3H), 1.26 – 1.23 (m, 2H), 1.08 (tdd, J = 12.2, 9.9, 4.1 Hz, 1H). LC-MS (ESI, m/z): $C_{28}H_{32}N_4O_2$, [M+H]+=457.2619.

**5-(3-(3-fluoropyrrolidin-1-yl)azetidin-1-yl)-2-methyl-N-(1-(quinolin-4-yl)cyclopropyl)benzamide formate (34)**. White solid, 13% yield. $^{1}$H NMR (600 MHz, DMSO-$d_6$) δ 9.15 (s, 1H), 8.87 (d, J = 4.4 Hz, 1H), 8.66 (dd, J = 8.5, 1.3 Hz, 1H), 8.05 (d, J = 8.4 Hz, 1H), 7.76 (ddd, J = 8.3, 6.6, 1.4 Hz, 1H), 7.69 (d, J = 4.4 Hz, 1H), 7.65 (ddd, J = 8.2, 6.7, 1.3 Hz, 1H), 6.91 (d, J = 8.2 Hz, 1H), 6.34 (dd, J = 8.2, 2.5 Hz, 1H), 6.12 (d, J = 2.5 Hz, 1H), 5.20 (dt, J = 55.4, 5.7 Hz, 1H), 3.80 (t, J = 7.1 Hz, 2H), 3.54 (td, J = 7.8, 5.2 Hz, 2H), 3.44 (ddd, J = 11.9, 6.7, 5.2 Hz, 1H), 3.30 (s, 1H), 2.83 – 2.72 (m, 2H), 2.63 (ddd, J = 32.0, 11.5, 4.9 Hz, 1H), 2.34 (td, J = 8.4, 5.8 Hz, 1H), 2.17 – 2.04 (m, 1H), 1.87 (s, 3H), 1.36 (q, J = 5.0 Hz, 2H), 1.25 – 1.23 (m, 2H). LC-MS (ESI, m/z): $C_{27}H_{27}FN_4O$, [M+H]+=445.2406.

**2-methyl-5-(3-(4-methylpiperazin-1-yl)azetidin-1-yl)-N-(1-(quinolin-4-yl)cyclopropyl)benzamide formate (35)**. White solid, 43% yield. $^{1}$H NMR (600 MHz, DMSO-$d_6$) δ 9.14 (s, 1H), 8.87 (d, J = 4.3 Hz, 1H), 8.66 (dd, J = 8.5, 1.4 Hz, 1H), 8.19 (s, 1H), 8.05 (dd, J = 8.5, 1.3 Hz, 1H), 7.76 (ddd, J = 8.3, 6.8, 1.4 Hz, 1H), 7.69 (d, J = 4.4 Hz, 1H), 7.65 (ddd, J = 8.3, 6.8, 1.3 Hz, 1H), 6.91 (d, J = 8.2 Hz, 1H), 6.34 (dd, J = 8.2, 2.5 Hz, 1H), 6.13 (d, J = 2.5 Hz, 1H), 3.81 (t, J = 7.0 Hz, 2H), 3.44 (dd, J = 7.4, 5.5 Hz, 2H), 3.19 (p, J = 6.1 Hz, 1H), 2.20 (s, 11H), 1.87 (s, 3H), 1.38 – 1.34 (m, 2H), 1.26 – 1.23 (m, 2H). LC-MS (ESI, m/z): $C_{28}H_{33}N_5O$, [M+H]+=456.2796.

**5-(4-(3-hydroxypyrrolidin-1-yl)piperidin-1-yl)-2-methyl-N-(1-(quinolin-4-yl)cyclopropyl)benzamide formate (36)**. White solid, 22% yield. $^{1}$H NMR (600 MHz, DMSO-$d_6$) δ 9.16 (s, 1H), 8.87 (d, J = 4.4 Hz, 1H), 8.67 (d, J = 8.4 Hz, 1H), 8.19 (s, 1H), 8.05 (d, J = 8.3 Hz, 1H), 7.76 (ddd, J = 8.3, 6.7, 1.4 Hz, 1H), 7.68 (d, J = 4.4 Hz, 1H), 7.67 – 7.62 (m, 1H), 6.94 (d, J = 8.4 Hz, 1H), 6.83 (dd, J = 8.4, 2.7 Hz, 1H), 6.60 (d, J = 2.7 Hz, 1H), 4.18 (q, J = 4.8, 3.5 Hz, 1H), 3.48 (d, J = 12.5 Hz, 2H), 2.78 (dd, J = 9.9, 6.0 Hz, 1H), 2.68 (d, J = 8.0 Hz, 1H), 2.64 – 2.56 (m, 2H), 2.42 (d, J = 9.8 Hz, 1H), 2.16 (s, 1H), 1.97 (ddt, J = 28.1, 14.3, 7.4 Hz, 2H), 1.89 (s,

2H), 1.85 (d, J = 12.6 Hz, 2H), 1.56 (d, J = 14.9 Hz, 1H), 1.43 (q, J = 10.4, 9.0 Hz, 2H), 1.37 (d, J = 2.2 Hz, 1H), 1.25 – 1.22 (m, 3H). LC-MS (ESI, m/z): $C_{29}H_{34}N_4O_2$, [M+H]+=471.2766.

**Procedure and characterization data for the synthesis of compound 19**

A mixture of 5-bromo-2-methyl-N-(1-(quinolin-4-yl)cyclopropyl)benzamide (1 equiv., 0.1 mmol, 38.1 mg), 1-methyl-4-(4,4,5,5-tetramethyl-1,3,2-dioxaborolan-2-yl)-1,2,3,6-tetrahydropyridine(1.2 equiv., 0.12 mmol, 37.1 mg), $Cs_2CO_3$ (2 equiv., 0.2 mmol, 65.2 mg), Pd2(dba)3 (0.1 equiv., 0.01 mmol, 9.1 mg), XPhos (0.2 eq., 0.02 mmol, 9.5 mg) and dioxane (1.5 mL) was added to a sealed tube. The mixture was stirred at 120 °C for 22 hours under an inert atmosphere. After that, water was added and the mixture was extracted with ethyl acetate twice. The organic phase was dried over $Na_2SO_4$ and concentrated. The residue was purified by preparative liquid chromatography. Lyophilization of the purified product afforded 16 mg desired solid.

**2-methyl-5-(1-methyl-1,2,3,6-tetrahydropyridin-4-yl)-N-(1-(quinolin-4-yl)cyclopropyl)benzamide formate (19)**. White solid, 56% yield. $^1$H NMR (600 MHz, DMSO-$d_6$) δ 9.26 (s, 1H), 8.87 (d, J = 4.4 Hz, 1H), 8.66 (dd, J = 8.4, 1.4 Hz, 1H), 8.17 (s, 1H), 8.05 (dd, J = 8.5, 1.3 Hz, 1H), 7.77 (ddd, J = 8.3, 6.8, 1.4 Hz, 1H), 7.70 (d, J = 4.4 Hz, 1H), 7.66 (ddd, J = 8.3, 6.8, 1.3 Hz, 1H), 7.32 (dd, J = 7.9, 2.1 Hz, 1H), 7.14 – 7.06 (m, 2H), 6.07 (s, 1H), 3.01 (d, J = 3.1 Hz, 2H), 2.56 (t, J = 5.7 Hz, 2H), 2.39 (tt, J = 3.7, 2.1 Hz, 2H), 2.28 (s, 3H), 1.99 (s, 3H), 1.42 – 1.35 (m, 2H), 1.27 – 1.25 (m, 2H). LC-MS (ESI, m/z): $C_{26}H_{27}N_3O$, [M+H]+=398.2232.

**Configuration of Tree-Invent running**

The configuration file can be divided into five components, namely “model”, “train”, “system”, “sample_constrain”, “rl”. The “model”, “train” and “system” settings are extracted from the hyper-parameters of training for the prior model. The initial learning rate “initlr” was set to 0.00005. The parameter “batchsize” was set to 20, which means that 20 molecules will be sampled per batch. In the component of “sample_constrain”, The nodes that must be generated are explicitly defined. The parameter “specific_nodefile” stores the structure of the pre-defined node in pickle file, which is set as the starting node 0. Atoms indexed in the “saturation_atomid_list” are not allowed for node growth. “node_conn” presents the atom in the pre-defined node for connection with node 1. “$atom_number” is the variable for setting the number of heavy atoms contained in node 1. In this study, for 5,6-fused and 6,6-fused ring, the variable is set to 9 and 10 respectively. The component “rl” is settings for reinforcement learning. Only docking score is selected as the scoring component, where the scoring weight is set to 1. The subcomponent “docking” provides the settings of molecular docking for generate docking scores.

```
{
    "model": {
        "device": "cuda",
        "mlp1_hidden_dim": 800,
        "mlp2_hidden_dim": 800
    },
    "train": {
        "batchsize": 20,
        "initlr": 0.00005
    },
    "system": {
        "max_atoms": 46,
        "max_cliques": 42,
        "max_rings": 8,
        "max_ring_size": 34,
        "max_ring_states": 62,
        "max_node_add_states": 42,
        "max_node_connect_states": 42,
        "ring_cover_rate": 0.97
    },
    "sample_constrain": {
        "max_node_steps": 4,
        "max_ring_nodes": 100,
        "temp": 1,
        "ring_check_mode": "easy",
        "constrain_step_dict": {
            "0": {
                "node add": {
                    "specific_nodefile": "specific_nodefile_0.pickle"
                },
                "node conn": {
                    "saturation_atomid_list": [
                        1,
                        2,
                        3,
                        4,
                        5,
                        6,
                        7,
```

```
                        8,
                        9,
                        10,
                        11,
                        12,
                        13,
                        14,
                        15,
                        16,
                        17,
                        18,
                        19
                    ]
                }
            },
            "1": {
                "node add": {
                    "max_ring_num_per_node": 2,
                    "min_ring_num_per_node": 2,
                    "max_aromatic_rings": 2,
                    "min_aromatic_rings": 1,
                    "max_heavy_atoms": $atom_number,
                    "min_heavy_atoms": $atom_number,
                    "force_step": true
                },
                "node conn": {
                    "constrain_connect_node_id": [
                        0
                    ]
                }
            }
        }
    "rl": {
        "acc_steps": 1,
        "temp_scheduler": "linear",
        "temp_range": [
            1.0,
            1.2
        ],
        "score_components": [
            "dockscore"
        ],
        "score_weights": [
            1
        ]
    },
    "docking": {
        "backend": "Glide",
        "dock_input_path": "$TreeInvent_path/$dock_input_path/",
        "dockstream_root_path": "$DockStream_path",
        "low_threshold": -8,
        "high_threshold": -5,
        "k": 0.3,
        "ncores": 8,
        "nposes": 3,
        "box_size": 20,
```

```
        "grid_path": "$glide-grid-file"
    }
}
```

### Supplementary Figures and Tables

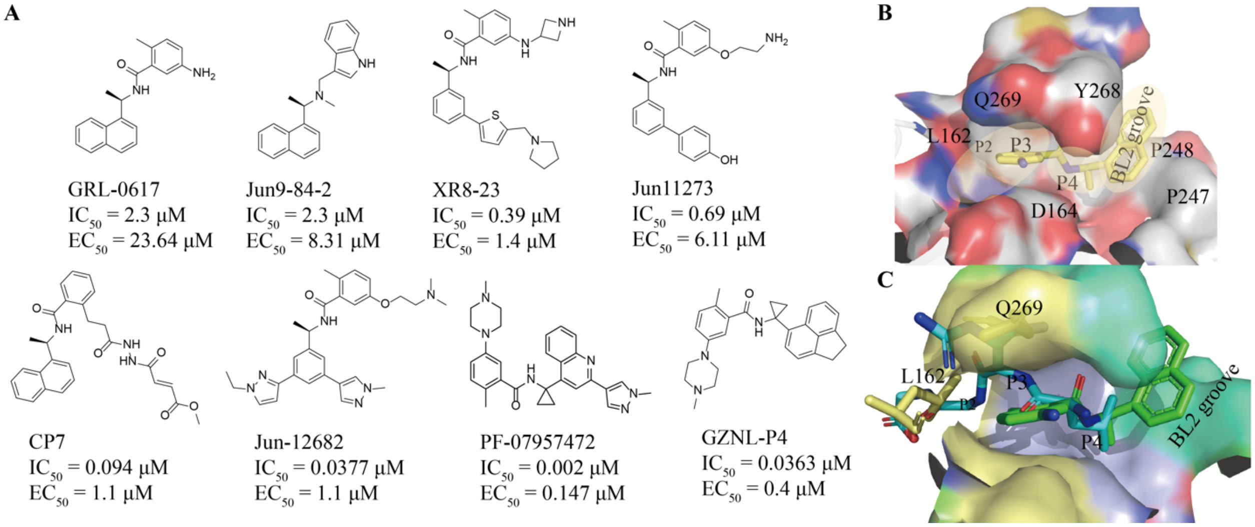

**Fig. S1. Structure and binding mode of available $PL^{pro}$ inhibitors.** A) Structure of some known SARS-Cov-2 $PL^{pro}$ inhibitors. $IC_{50}$ and $EC_{50}$ are the enzymatic and antiviral activity for wild-type $PL^{pro}$, respectively. B) Residues of $PL^{pro}$ involved in the interaction with **GRL0617 (**PDB ID: 7JRN**)**. The binding pocket was shown as a protein surface and colored by element types. Ligand was shown as yellow sticks. Dashed circles represent the sub-pockets. C) Overlap between **GRL0617** and substrate peptide (LRGG) in the active pocket of $PL^{pro}$. The substrate peptide is extracted from the co-crystal structure of SARS-COV $PL^{pro}$ with ubiquitin (PDB ID: 4M0W)[40]. The BL2 groove, P4 and P3 sub-pockets are shown as green, blue and yellow surface respectively. Green and cyan sticks refer to the **GRL0617** and substrate peptide (sequence LRGG). The residues L162 and Q269 are shown as yellow sticks.

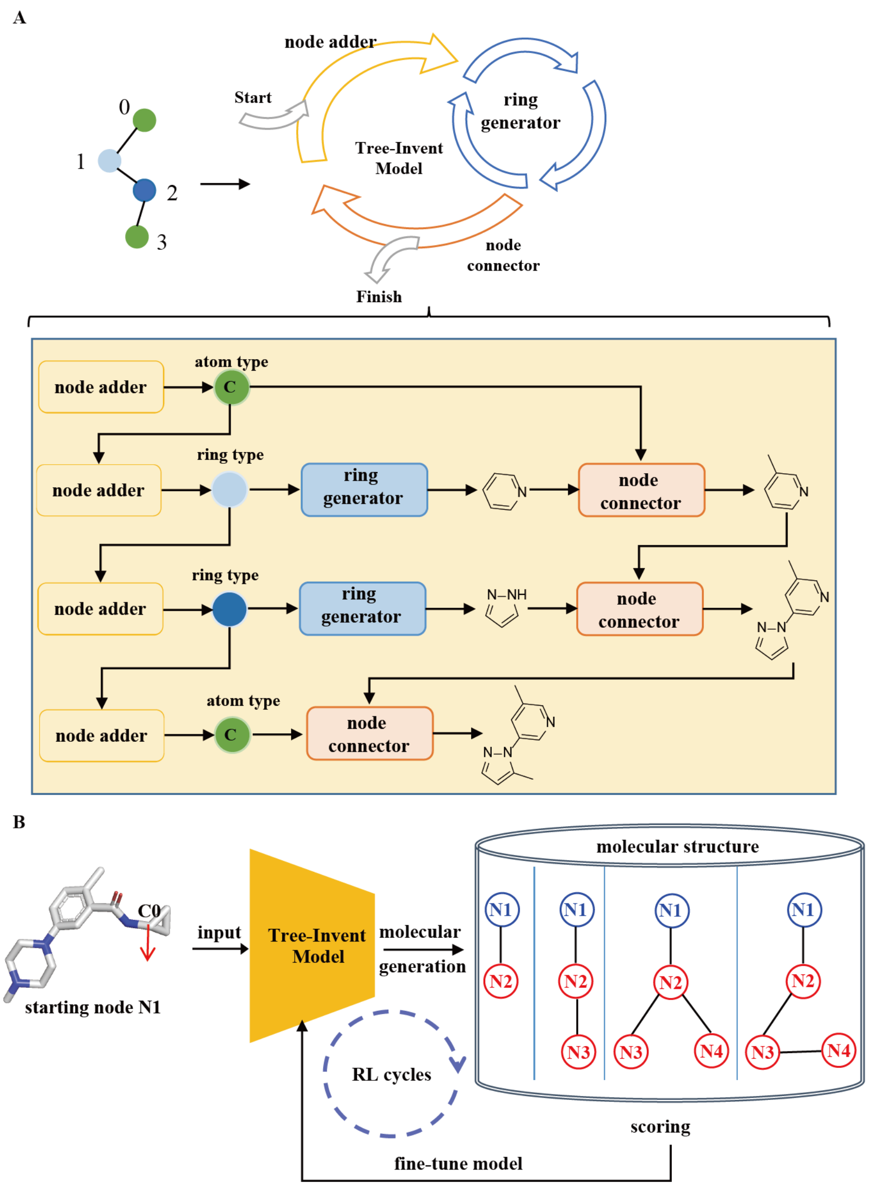

**Fig. S2. Molecular generation method with Tree-Invent**. A) Scheme of molecular generation under a topology constraint. B) Basic workflow for molecular generation based on the substructure of GZNL-P4 using the Tree-Invent model with RL strategy.

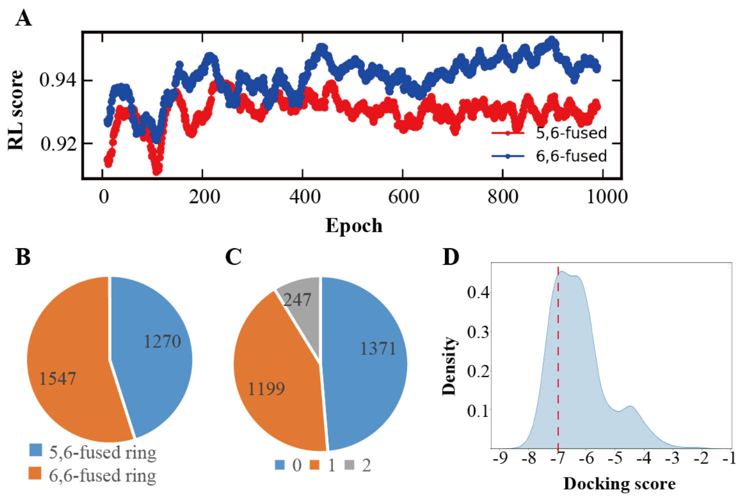

**Fig. S3. Statistics for molecular generation using Tree-Invent.** A) Learning curves of reinforcement learning. B) Number of molecules with 5,6-fused and 6,6-fused ring at $R^2$ position. C) Number of molecules with zero, one and two substitutes on the ring of the $R^2$ group. D) Distribution of docking score of all generated valid and unique molecules using the kernel density estimation method. The X axis is the docking score, while the Y axis is the probability density under this estimation.

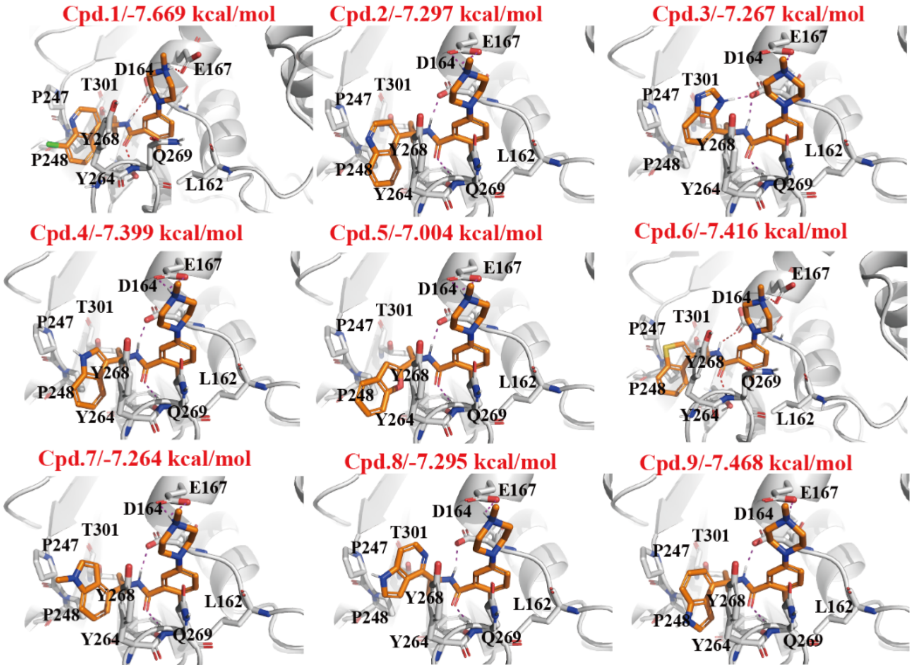

**Fig. S4. Docking pose and docking score of Compound 1-9 with PL$^{pro}$.** The compounds were docked to the co-crystal structure of PL$^{pro}$ with GRL0617 (RCSB PDB code: 7JRN). The protein secondary structure is shown as gray cartoons. Ligand is shown as orange sticks. Red dashes represent hydrogen bond and salt bridge interactions.

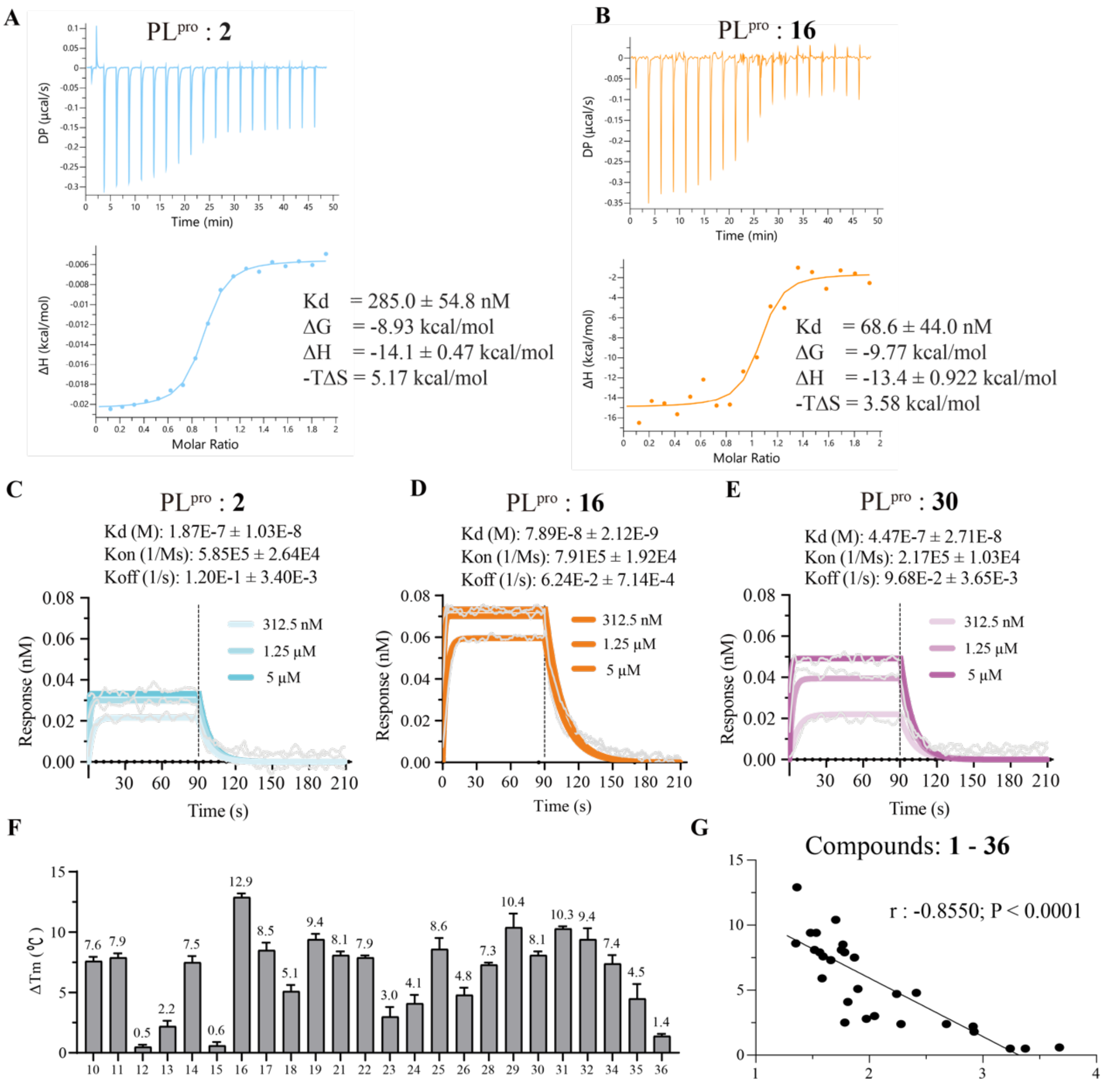

**Fig. S5. Binding affinity determined by ITC, BLI and DSF.** A-B) Isothermal titration calorimetry (ITC) binding curve for the interactions of SARS-CoV-2 PL$^{pro}$ with compound 2 (A) and 16 (B). C-E) Bio-layer interferometry (BLI) binding curve for the interactions of SARS-CoV-2 PL$^{pro}$ with compound 2 (C), 16 (D) and 30 (E). F) Differential scanning fluorimetry (DSF) assay analysis of SARS-CoV-2 PL$^{pro}$ with compounds optimized from compound **2**. Data are presented as mean ± s.d. of three technical replicates. G) The correlation analysis of designed compounds between the inhibition activity and delta Tm determined by DSF.

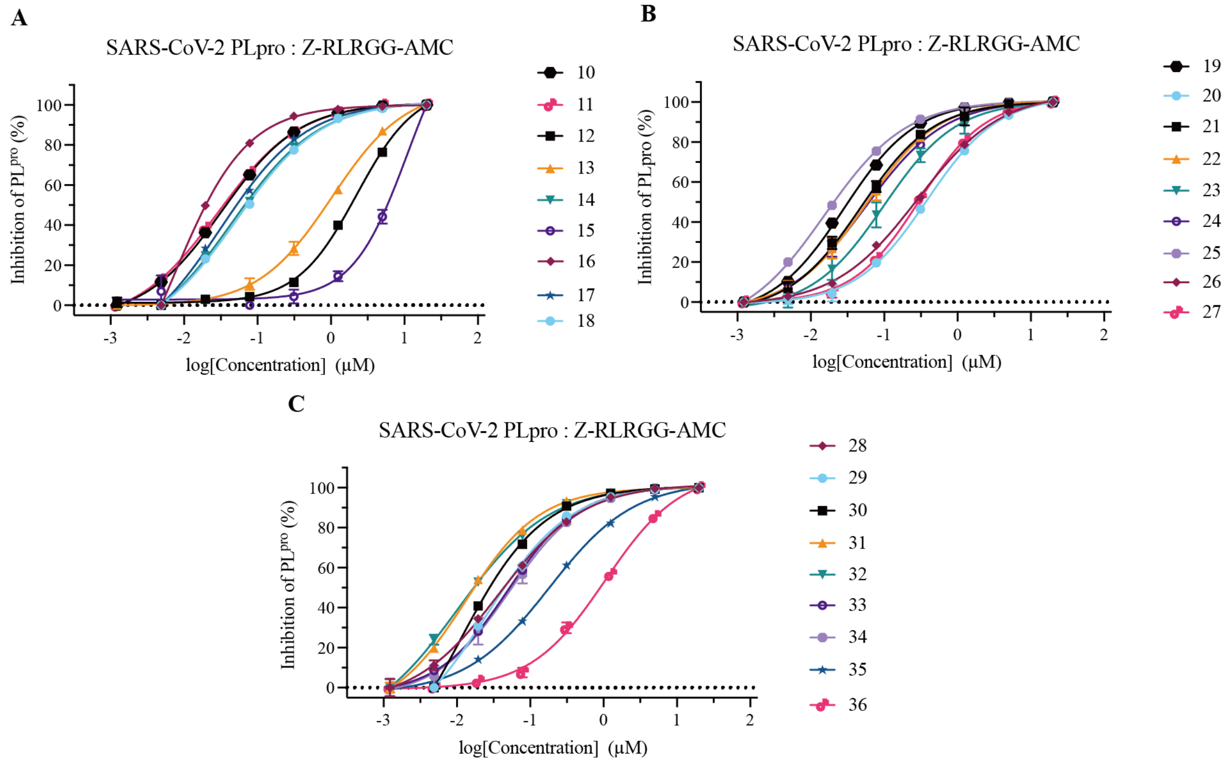

**Fig. S6. Dose dependent inhibition rate of compounds 10-18 (A), 19-27 (B) and 28-36 (C) at the enzymatic level.** Data are presented as mean ± s.d. of three technical replicates.

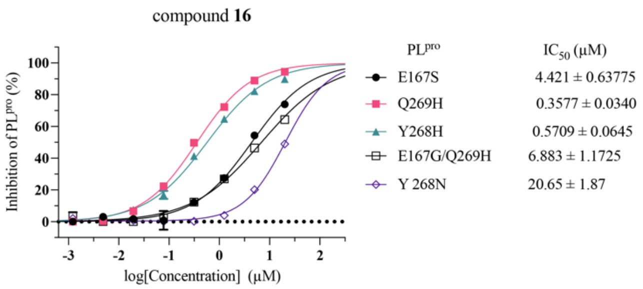

**Fig. S7. The enzymatic inhibition activity of compound 16 for PL$^{pro}$ mutants**. Data are presented as the mean of two technical replicates.

### Table S1. X-ray data collection and refinement statistics

| PL$^{pro}$ in complex with ligands | | | |
|---|---|---|---|
| | GZNL-2002 | GZNL-2016 | GZNL-2030 |
| **Data collection** | SSRF-BL02U1 | SSRF-BL02U1 | SSRF-BL02U1 |
| **Space group** | P $4_1$ 2 2 | P 1 $2_1$ 1 | P 1 $2_1$ 1 |
| **Cell dimensions** | | | |
| ***a*, *b*, *c* (Å)** | 95.18, 95.18, 232.35 | 46.57, 144.26, 59.74 | 46.30, 144.61, 60.12 |
| ***α*, *β*, *γ* (°)** | 90, 90, 90 | 90.00, 98.98, 90.00 | 90.00, 99.48, 90.00 |
| **Resolution** | 73.63 – 2.80 (2.95 – 2.80) | 54.61 – 2.30 (2.38 – 2.30) | 72.31 – 2.70 (2.83 – 2.70) |
| $R_{pim}$ | 0.051(0.166) | 0.057(0.390) | 0.094(0.370) |
| **I / σ(I)** | 8.9(3.4) | 9.4(2.5) | 5.8(2.2) |
| ***CC*1/2 in highest shell** | 0.935 | 0.995 | 0.985 |
| **Completeness (%)** | 95.65 (98.0) | 99.9 (100) | 89.9 (98.0) |
| **Redundancy** | 3.2 (3.1) | 6.9 (7.0) | 2.6 (2.6) |
| **Refinement** | | | |
| **Resolution (Å)** | 2.80 | 2.30 | 2.70 |
| **No. of unique reflections** | 26015 | 34510 | 19237 |
| $R_{work}/R_{free}$ **(%)** | 20.9/25.7 | 21.96/24.69 | 22.22/28.29 |
| **No. of atoms** | | | |
| **Protein** | 4961 | 4947 | 4908 |
| **Water** | 12 | 50 | 1 |
| ***B*-factors** | | | |
| **Protein** | 57.46 | 46.70 | 50.24 |
| **Ligand** | 40.35 | 40.65 | 44.22 |
| **Water** | 44.45 | 37.86 | 46.38 |
| **RMSD** | | | |
| **Bond lengths (Å)** | 0.010 | 0.012 | 0.015 |
| **Bond angles (°)** | 1.166 | 1.854 | 1.521 |
| **Ramachandran favored (%)** | 93.06 | 96.41 | 96.05 |
| **Ramachandran outliers (%)** | 0.32 | 0 | 0 |
| **PDB accession code** | 8ZSE | 21MS | 21MU |

****Values in parentheses indicate highest resolution shell.***
